# Control of Inflammatory Response by Tissue Microenvironment

**DOI:** 10.1101/2024.05.10.592432

**Authors:** Zhongyang Wu, Scott D. Pope, Nasiha S. Ahmed, Diana L. Leung, Stephanie Hajjar, Qiuyu Yue, Diya M. Anand, Elizabeth B. Kopp, Daniel Okin, Weiyi Ma, Jonathan C. Kagan, Diana C. Hargreaves, Ruslan Medzhitov, Xu Zhou

## Abstract

Inflammation is an essential defense response but operates at the cost of normal tissue functions. Whether and how the negative impact of inflammation is monitored remains largely unknown. Acidification of the tissue microenvironment is associated with inflammation. Here we investigated whether macrophages sense tissue acidification to adjust inflammatory responses. We found that acidic pH restructured the inflammatory response of macrophages in a gene-specific manner. We identified mammalian BRD4 as a novel intracellular pH sensor. Acidic pH disrupts the transcription condensates containing BRD4 and MED1, via histidine-enriched intrinsically disordered regions. Crucially, a decrease in macrophage intracellular pH is necessary and sufficient to regulate transcriptional condensates *in vitro* and *in vivo*, acting as negative feedback to regulate the inflammatory response. Collectively, these findings uncovered a pH-dependent switch in transcriptional condensates that enables environment-dependent control of inflammation, with a broader implication for calibrating the magnitude and quality of inflammation by the inflammatory cost.

**Highlights:**

- Acidic pH regulates a switch-like gene-specific inflammatory response in macrophages
- Acidic pH impacts chromatin remodeling and transcription circuits to control inflammatory programs
- BRD4 transcriptional condensates are regulated by intracellular pH via the histidine-enriched intrinsically disordered region
- Tissue inflammation decreases intracellular pH and disrupts BRD4 condensates as a negative feedback

## Introduction

Inflammation is crucial for maintaining homeostasis and defending the integrity of tissues and organs. Yet, excessive inflammatory responses can result in significant tissue damage.^1^ Much of our knowledge in the protective and pathological roles of inflammation relates to how inflammatory responses impact organ functions. By contrast, less is known about how the state of a given tissue impacts the inflammatory response. The magnitude, duration and specific impact of an inflammatory response must be calibrated to both the presence of inflammatory triggers and the extent of pathological outcome.^2^

The maintenance of a stable pH range is a universal feature of tissue homeostasis. Different tissues and organs maintain an interstitial environment within specific pH range, whereas these pH levels are often perturbed during inflammation.^3^ For instance, blood pH is tightly regulated between 7.35 and 7.45 through respiration and renal compensation.^4^ In patients with sepsis, severe acidosis indicates a poor prognosis for survival.^5,6^ In the brain, the cerebrospinal fluid maintains a mildly acidic pH ∼7.3; however, ischemic injury can lower this to pH 6.6.^7^ The pH of the paracortical zone of lymph nodes is sustained between 6.3 and 7.1, and becomes even more acidic upon infections.^8^ In addition, aberrant microenvironment of solid tumors frequently exhibit acidic pH due to heightened metabolic activity, hypoxia, and active proton extrusion by cancer cells.^9^ These deviations from normal pH levels can indicate an emergent state in affected tissues.

In mammalian cells, pH levels can be monitored both extracellularly and intracellularly. On the cell surface, a broad range of pH values can be detected by a variety of extracellular sensors, including G-protein coupled receptors (GPR4, GPR65, GPR68), acid sensing ion channels (ASIC1, ASIC2, ASIC3, ASIC4) and transient receptor potential cation channel subfamily V member 1 (TRPV1).^4^ These extracellular pH sensors are activated at the pH range of 4 to 8,^10,11^ regulating various cellular and physiological parameters, from blood pH and lipid metabolism to pain sensations during exercise.^12–14^ In contrast, the mechanisms intracellular pH sensing and the subsequent regulation of cellular responses are less understood. The Hypoxia-Induced Factors, HIF1 alpha and HIF2 alpha, can be activated by acidic conditions independently of hypoxia.^15,16^ In the context of cancers, transcription factors SMAD5 and Sterol Regulatory Element-Binding Protein 2 (SREBP2) have been implicated in cellular responses to intracellular pH in cancer cell lines.^17,18^ It remains to be determined whether and how pH-sensing mechanisms specifically regulate cellular activities in the context of inflammation. Despite this lack of understanding, acidic pH is generally considered suppressive to cell activation and proliferation. Change in pH may influence survival, differentiation, migration and cellular metabolism in a cell-type specific manner.^3^ In considering immune cells that may sense changes in pH and influence tissue activity, we focused on macrophages given their role as tissue sentinels.^19,20^ In response to cues associated with microbial infections, macrophages regulate the expression of thousands of genes important for tissue homeostasis, innate host defense, antiviral response, and coordination with adaptive immune system.^21^ We sought to understand how detecting pH changes in macrophages regulates the inflammatory response to pathogens.

Recent studies revealed that proteins can form biomolecular condensates of fundamental roles in cellular organization, signaling, stress response and gene regulation.^22–27^ In particular, critical factors involved in gene transcription tend to concentrate in distinct and dynamic nuclear foci referred to as transcriptional “hubs” or “condensates”. These mechanisms represent crucial biochemical controls of gene expression, transcription burst and compartmentalization of opposing regulators.^28–33^ The proteins identified within these foci, including BRD4, MED1, Pol II, YAP/TAZ, TBP, P300, pTEFb, typically contain intrinsically disordered regions (IDRs) that favor weak and multivalent interactions.^32–39^ Specifically, BRD4 is a member of the bromodomain and extraterminal (BET) protein family that recognizes histone lysine acetylation associated with active transcription.^40,41^ It interacts and recruits the mediator complex, pTEFb and other transcription machinery, as an essential regulator of Pol II-dependent transcription.^42,43^ In macrophages and in mice, inhibition of BRD4 reduces inflammatory response and prevents lethality in severe sepsis,^44^ suggesting that BRD4 function is vital for the activation of inflammatory response. Despite the increasing examples of transcriptional condensates in gene regulation, control of transcriptional condensate formation and their roles in the immune system are not well understood.

In this study, we explored the hypothesis that macrophages use the detection of intracellular pH deviation as a guide to control inflammatory responses (Figure 1A). Using the Toll-like receptor 4 (TLR4) ligand bacterial lipopolysaccharide (LPS), we found that changes in pH do not impact TLR4 signal transduction *per se*, but strongly impact the spectrum and the extend of inflammatory genes activated by TLR4. We identified BRD4 as a novel intracellular pH sensor. The BRD4-containing transcription condensates are pH-sensitive, regulated by the evolutionary conserved histidine-enriched IDR. Moreover, the inflammatory activation of macrophages triggers intracellular acidification and alters transcriptional condensates *in vitro* and *in vivo*. Thus, this work reveals a new regulatory mechanism of inflammation, where transcriptional condensates integrate extracellular and intracellular pH via BRD4 to elicit gene-specific inflammatory response based on microenvironment. Sensing pH deviation by the immune system may act as a feedback mechanism to balance the protective benefits of immune response against their potentially pathological impact.

**Figure 1.**
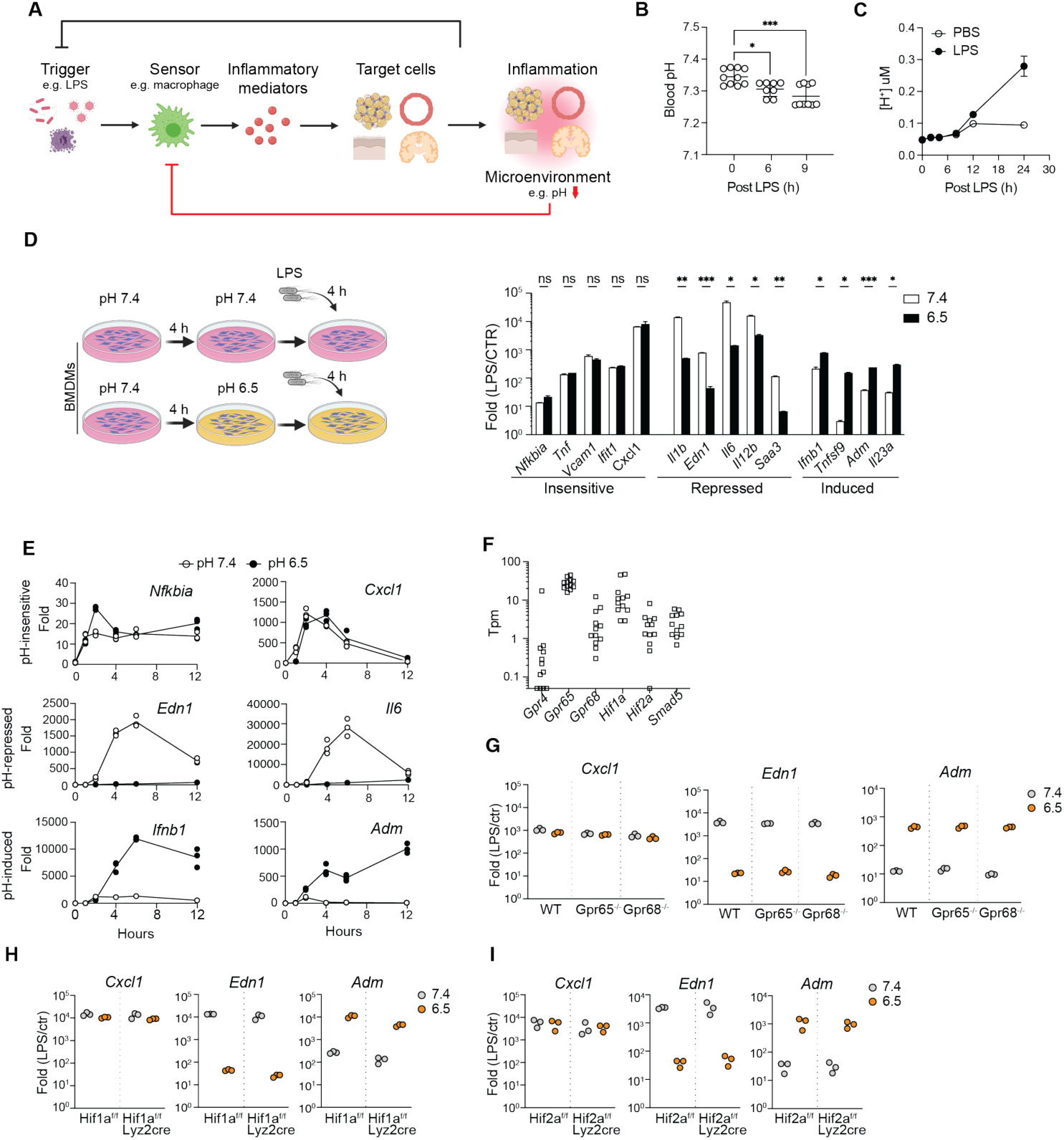
Acidic pH regulates gene-specific inflammatory response in macrophages. (A) A diagram of an inflammatory circuit with pH-sensing feedback to the innate sensors. (B) Blood pH in mice challenged with 10 mg/kg LPS intraperitoneally. One-way ANOVA, Dunnett test for multiple comparisons, Mean +/-STD. (**C**) Extracellular acidification of BMDMs stimulated with 100 ng/mL LPS *in vitro*. Mean+/-STD. (**D**) Fold activation of inflammatory genes in BMDMs after 4 hours 10 ng/mL LPS at pH 7.4 or 6.5, normalized to unstimulated conditions respectively. Mean+/-standard deviation (STD). Unpaired t-test, Holm-Sidak’s test for multiple comparisons. (**E**) Time course analysis of gene activation in BMDMs stimulated with 10 ng/mL LPS at pH 7.4 and 6.5. (**F**) Expression of known pH sensors in macrophages, including BMDMs, monocytes, tissue resident macrophages in peritoneal, liver, lung, small and large intestines (ref: Lavin 2014). (**G-I**) Fold changes of selected genes in WT, *Gpr65-/-* and *Gpr68*-/-BMDMs (G), *Hif1a*-/-BMDMs (H) and *Hif2a*-/-BMDMs (I). In (B) and (D), *p<0.05, **p<0.01, ***p<0.001.

## Results

### Acidic pH modulates the gene-specific inflammatory response

In a murine model of acute inflammation, intraperitoneal (i.p.) injection of a sub-lethal dose of LPS in wild-type mice (WT) triggered a TLR4-dependent systemic inflammatory response. Consequently, the blood pH decreased significantly after 6 hours of injection (Figure 1B) and severe acidosis (< pH 6.5) was reported in various tissues at 24 hours,^45^ underscoring the link between acidic pH levels and acute inflammatory response. *In vitro*, WT bone-marrow derived macrophages (BMDMs) acidified the extracellular environment to pH 6.5 at 24 hours after LPS stimulation (Figure 1C and S1A), in the presence of a physiological level buffer system (23.8 mM bicarbonate). Notably, this pH level did not adversely affect the viability of BMDMs (Figure S1B) nor impede TLR4 internalization triggered by LPS exposure (Figure S1C). Therefore, we conditioned BMDMs at pH 6.5 to investigate the influence of acidic pH on the inflammatory responses of macrophages.

The transcriptional response to LPS in macrophage is regulated by TLR4 signaling pathways, epigenetic mechanisms, and both feedback and feedforward genetic circuits.^21,46–49^ The transcriptionally induced genes can be categorized into primary response genes (PRGs) and secondary response genes (SRGs) depending on the reliance of the latter on newly synthesized proteins.^21^ PRGs can be further subdivided into early and late response groups based on activation kinetics. To evaluate the overall impact on the inflammatory response, we analyzed expression of early PRGs (*Cxcl1*, *Il1b, Nfkbia*, *Tnf*, *Tnfsf9*), late PRGs (*Ifnb1*) and secondary response genes (*Ifit1*, *Il6*, *Il12b*, *Saa3*) at 4 hours post LPS exposure (Supplemental table 1).^50^ We found that conditioning BMDMs at pH 6.5 revealed a gene-specific impact on inflammatory genes that did not correlate with the classification of PRGs and SRGs (Figure 1D). The transcription of *Cxcl1, Ifit1, Nfkiba, Tnf* and *Vcam1* was relatively insensitive to pH, whereas *Edn1*, *Il1b*, *Il6*, *Il12b* and *Saa3* were significantly repressed at pH 6.5. Remarkably, the induction of *Ifnb1*, *Tnfsf9*, *Adm* and *Il23a* was greatly enhanced, showing an average of 20-fold increase at pH 6.5. Varying experimental conditions, including concentrations of LPS (10 to 1000 ng/mL) and duration of pH conditioning (4-12 hours) demonstrated a similar pattern of gene-specific regulation by acidic pH (Figure S1D, E). Furthermore, time-course analyses of gene expression displayed a switch-like pattern among pH-regulated genes (Figure 1E). Notably, *Edn1* and *Adm*, which encode endothelin-1 and adrenomedullin respectively, exhibit differential activation at pH 7.4 and 6.5. These two peptides have antagonistic effect on blood vessels through vasoconstriction and vasodilation, respectively.^51^ The gene-specific sensitivity to the acidic environment and the regulation of opposing physiological functions suggest that macrophages may orchestrate a qualitatively different type of inflammatory response based on microenvironment, rather than merely adjusting the magnitude of immune activation.

### Known pH sensors cannot account for the pH-dependent inflammatory response

Among the known pH sensors, *Gpr65*, *Gpr68*, *Hif1a*, *Hif2a* are abundantly expressed in BMDMs and tissue resident macrophages (Figure 1F, S1F).^52^ We investigated pH-dependent inflammatory responses in BMDMs differentiated from *Gpr65*^-/-^, *Gpr68*^-/-^, *Hif1a^flox/flox^ lyz2^Cre^* and *Hif2a^flox/flox^ lyz2^Cre^* mice *in vitro*. However, our results indicated that neither *Gpr65*, *Gpr68*, *Hif1a* nor *Hif2a* alone was essential for the observed gene-specific, pH-dependent regulation (Figure 1G-I). At pH 6.5, both GPR65 and GPR68 activate the production of cyclic AMP (cAMP).^10^ To exclude the possibility of receptor redundancy, we treated BMDMs with 100 μM dibutyryl-cAMP, a cell-permeable analog of cAMP, and observe no significant effect on pH-dependent gene expression (Figure S1G). These findings suggested that the observed response was independent of established pH sensing mechanisms. Although acidic pH may broadly affect biochemical reactions and protein interactions, the specific activation and repression of genes at pH 6.5 imply that a general interference with transcription was unlikely. To further test this, we treated BMDMs with camptothecin, a DNA topoisomerase inhibitor that broadly represses the inflammatory response of macrophages by inhibiting Pol II.^53^ Camptothecin inhibited the activation for pH-insensitive and pH-sensitive genes (Fig S1H). Thus, we hypothesized that a novel specific pH-sensing mechanism controls inflammatory gene expression.

### A deconvolution model reveals pH-dependent inflammatory programs

We compared bulk RNA-seq of BMDMs at pH 7.4, pH 7.4 LPS 4 h, pH 6.5, and pH 6.5 LPS 4 h (Figure 2A) to comprehensively characterize the pH-dependent and pH-independent inflammatory response. Applying a stringent threshold (Fold change > 3 and q-value < 0.05), we identified 522 genes uniquely induced at pH 7.4 and 85 genes uniquely induced at pH 6.5, underscoring the gene-specific inflammatory response by acidic pH. However, among 308 genes qualified as induced at both pH 7.4 and pH 6.5, many differed in their activation and expression quantitatively (Figure S2A). To gain a quantitative understanding of how the inflammatory response is regulated by acidic pH, we employed a linear deconvolution model originally developed to dissect regulatory interactions among transcription factors.^54,55^ This model conceptualizes that two signals, such as LPS stimulation and acidic pH, can control gene expression through three possible logics: LPS regulation independent of pH (LPS), pH regulation independent of LPS (pH), and the regulation dependent on the synergistic or antagonistic interactions between LPS and pH (INT). These 3 regulations (referred to as “expression components”) would describe the observed differential gene expression among experimental conditions (Figure 2B). Applying the linear deconvolution model allows integrating all four conditions simultaneously to assess how LPS, pH, and their interactions contribute to the expression of each gene (Figure S2B). An expression component close to 0 indicates a lack of regulation, while a positive or a negative value indicates activation or repression, or that LPS and acidic pH act synergistically or antagonistically.

**Figure 2.**
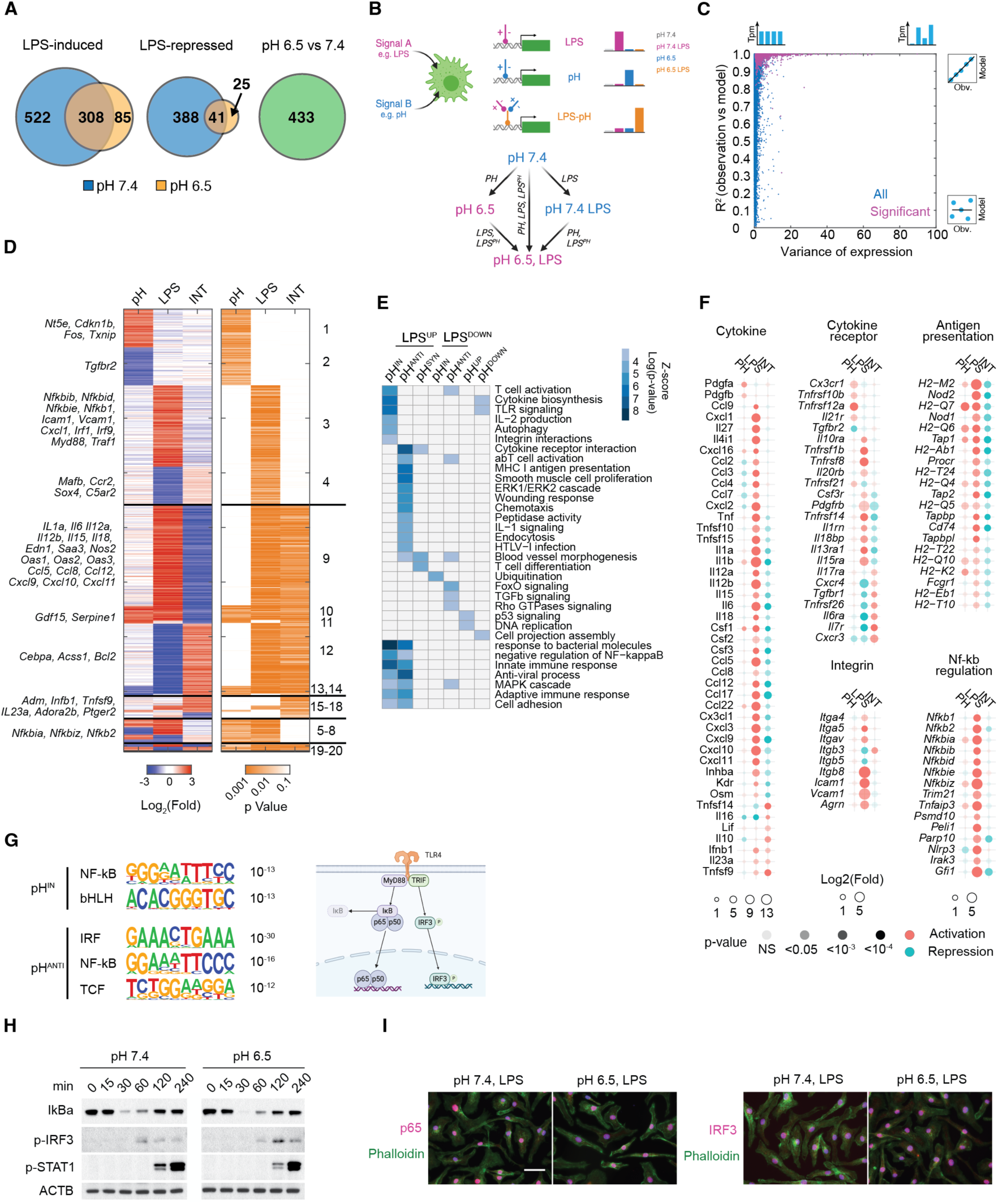
Deconvolution analysis revealed pH-dependent combinatorial control of inflammatory response. (**A**) Van diagrams of significantly regulated genes by LPS or acidic pH. Fc > 3 and q < 0.05. (**B**) Illustration of the linear deconvolution model to identify gene regulatory logics between LPS and acidic pH. (**C**) Evaluation of linear model fitting using expression variance and R-square for each gene. Blue marks all genes and magenta marks differentially expressed genes (FC > 3, q<0.05) in any pair of conditions. (**D**) Heatmap of inflammatory genes regulated by either LPS stimulation or acidic pH. Left, log2(fold) of model-inferred regulation by acidic pH alone (pH), LPS stimulation alone (LPS) and the interactions between acidic pH and LPS stimulation (LPS^PH^). Right, heatmap of p-values of each expression component, determined with a null hypothesis that the regulatory effect is less than 1.5-folds. Cluster groups (1-20) were determined based on expression component p-values. (**E**) Functional enrichment for LPS-induced or LPS-repressed genes that are pH-insensitive, pH-antagonistic or pH-synergistic. (**F**) Expression components of inflammatory cytokines, cytokine receptors, antigen presentation genes, integrins and regulators of NF-ΚB signaling. (**G**) Motif enrichment of LPS-induced genes that are pH-insensitive and pH-antagonistic. (**H**) Western blot of NF-ΚB, IRF3 and STAT1 signaling activation in BMDMs after LPS stimulation. (**I**) Immunofluorescence staining of p65 (pink) and IRF3 (pink) in BMDMs after LPS stimulation, 1 hour for p65 and 2 hours for IRF3.

Overall, more than 92% of the genes significantly regulated by pH or LPS (Figure S2C) were well captured using the 3-component linear model (R^2^ > 0.9) (Figure 2C). We identified 1620 genes with at least one significant expression component (p<0.05, null hypothesis component > 2-fold), forming 20 clusters with at least 5 genes each (Figure 2D, S2D, Supplemental table 2). Among genes regulated by LPS and pH independently, Cluster 1-4 represent independent activation or repression by either acidic pH or LPS, while Clusters 5-8 represent combinations of these independent regulations. For example, Cluster 3 includes *Nfkbib*, *Nfkbid*, *Nfkbie* and Cluster 5 includes *Nfkbia* and *Nkfbiz*, all belonging to the NF-ΚB signaling pathway. They are induced by LPS independent of pH and have differences in pH-dependent basal expression (Figure S2F). Cluster 9-14 include genes regulated antagonistically by pH and LPS (a positive LPS and a negative INT expression component), including inflammatory cytokines (*Il6, Il12a, Il12b, Il18*), chemokines (*Ccl5, Ccl8, Ccl12, Cxcl9, Cxcl10, Cxcl11*), acute phase proteins (*Oas1, Oas2, Oas3, Saa3*) and inflammatory effectors (*Edn1, Nos2*). At last, group 15-18 include genes synergistically regulated by LPS and acidic pH, such as *Adm, Ifnb1, Tnfsf9, Il23a* and *Adora2b*. To examine potential immune functions enriched in pH-dependent and independent responses, we merged the 20 clusters into groups characterized as pH-insensitive (non-significant INT), pH-antagonistic (opposite signs between LPS and INT) or pH-synergistic (both positive or both negative for LPS and INT) (Figure S2D). Overall, approximately 40% of LPS-regulated genes are pH-insensitive. Among pH-regulated genes, an antagonistic effect predominated in both LPS-induced and LPS-repressed responses (88% and 86% respectively, Figure S2E). Since LPS induced transcriptional program serves as a model system for innate inflammatory response, we focus on the three groups of LPS induced genes: pH-insensitive (pH^IN^), pH-repressed (pH^ANTI^) and pH-synergistic genes (pH^SYN^). pH^IN^ genes were uniquely enriched in TLR signaling, T cell activation, integrin interactions, and showed strongest enrichment in innate immune response and anti-microbial defense (Figure 2E). The majority of LPS-induced integrins and NF-ΚB regulators are regulated by LPS independent of pH (Figure 2F). On the contrary, pH^ANTI^ genes were uniquely enriched in MHC I presentation, IL-1 signaling, chemotaxis, and showed the strongest enrichment in cytokine receptor interactions, antiviral and adaptive immune response (Fig. 2E). The activation of most antigen presentation genes and cytokines were strongly repressed by acidic pH (Figure 2F). Interestingly, pH^SYN^ genes, although fewest in number, were enriched in blood vessel morphogenesis and T cell differentiation (Figure 2E). These analyses revealed that the activation of innate defense programs is relatively insensitive to pH, while the coordination and recruitment of other branches of the immune system seem to be dependent on the tissue microenvironment.

TLR4-induced genes are regulated by two signaling adaptors, MyD88 and TRIF (Figure 2G).^49^ MyD88 recruits IL-1 receptor-associated kinases (IRAKs) and TRAF6, initiating the activation of the transcriptional activator NF-ΚB. Conversely, TRIF facilitates TRAF3-dependent activation of TBK1, which phosphorylates the transcription factor IRF3 to regulate interferon beta and other interferon-induced genes. We found that the consensus binding motif of p65 (NF-ΚB) was enriched in both pH^IN^ and pH^ANTI^ genes, whereas the IRF3 motif was only enriched in pH^ANTI^ genes (Figure 2G). However, LPS-induced degradation of the NF-ΚB inhibitory regulator IΚBα, IRF3 phosphorylation as well as autocrine type-I Interferon (induced by both NF-ΚB and IRF3) signaling via STAT1 phosphorylation, all displayed comparable kinetics between pH 7.4 and pH 6.5 in BMDMs (Figure 2H). Furthermore, the nuclear localization of p65 and IRF3 was found comparable at both pH levels (Figure 2I). These findings suggest that pH-sensitive and pH-insensitive genes are differentially regulated at the level of transcriptional activation, rather than signal transduction.

### pH regulates gene expression at the chromatin level

IRF3 exhibited a delayed activation kinetics compared to NF-ΚB (Figure 2H). The observed enrichment of IRF3 motifs indicated possible difference in activation kinetics between pH^IN^ and pH^ANTI^ genes. At pH 7.4, we found that pH^IN^ genes reached full activation within 2 hours after LPS stimulation, whereas pH^ANTI^ genes showed minimal induction, only peaking at 4 hours (Figure 3A). Interestingly, pH^SYN^ genes demonstrated the most rapid and transient activation (Figure 3A, Figure S3A). The LPS-repressed genes also displayed a similar pH-dependent kinetics (Figure S3B). Therefore, acidic pH may act on a time-limiting step required for the activation of the pH^ANTI^ genes. LPS-induced SRGs typically have delayed activation due to the requirement for new protein synthesis,^50^ and are found to have a strong pH-dependence compared to PRGs in our data (Figure S3C). However, blocking protein synthesis in BMDMs with cycloheximide (CHX) did not affect pH^ANTI^ genes to the same extent as acidic pH (Figure 3B, S3D). On the other hand, the activation of pH^SYN^ genes was almost completely replicated by blocking protein synthesis (Figure 3B, S3D), indicating that an induced transcriptional repressor may be suppressed by acidic pH, thereby facilitating the elevated activation of pH^SYN^ genes.

**Figure 3.**
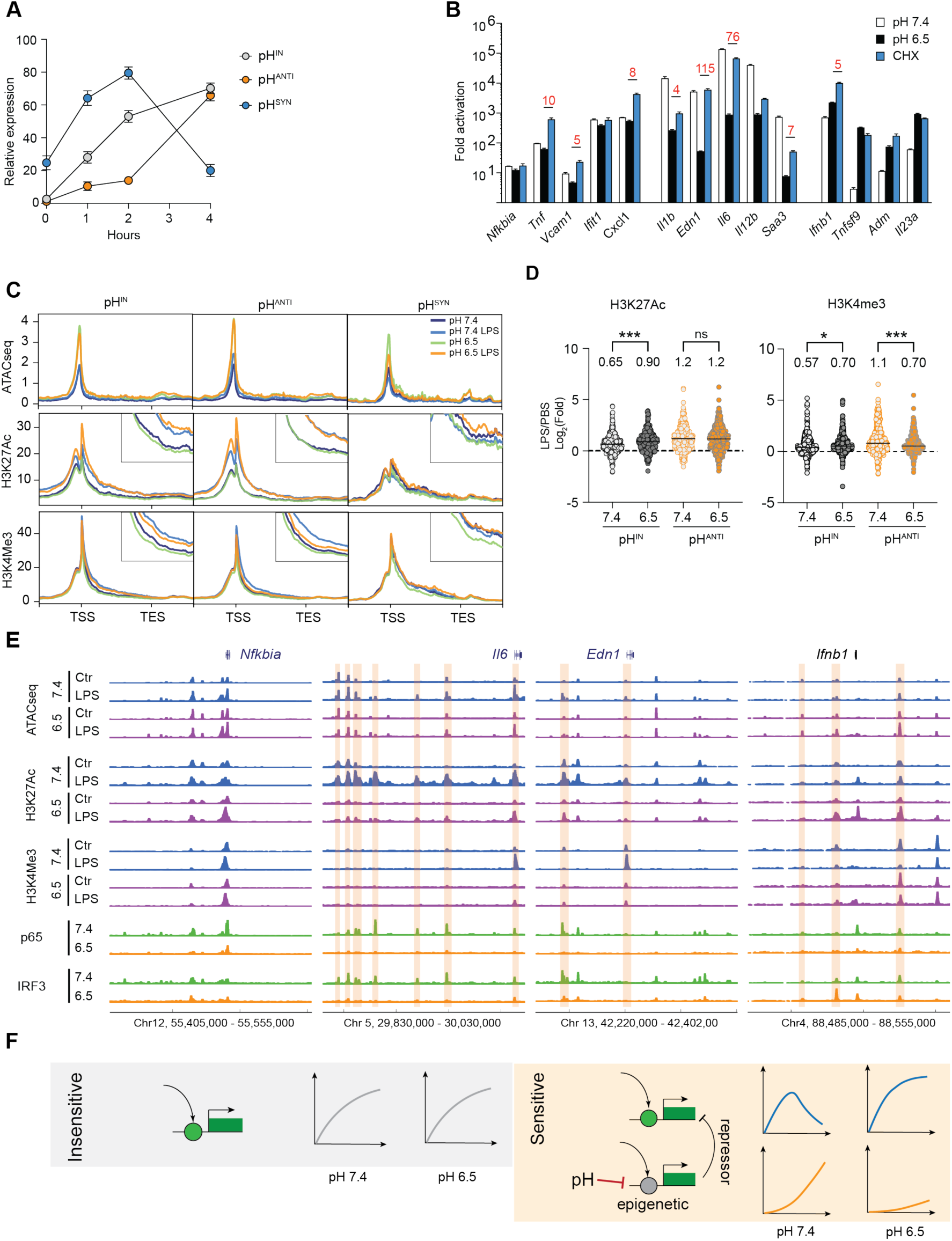
Transcription circuit and epigenetic control underlies environment-sensitive inflammatory response. (**A**) Expression kinetics of LPS-induced genes for top pH-insensitive (100), pH-antagonistic (100) and pH-synergistic groups (58). (**B**) Activation of inflammatory genes by LPS with 10 ng/mL for 4 hours, at pH 7.4, 6.5 or with 200 ng/mL cycloheximide (CHX). Fold change is calculated as LPS/untreated in each treatment condition. (**C**) Average profile of ATAC-seq and ChIP-seq of H3K27Ac and H3K4me3 of pH-regulated genes. Intersection image shows an enlarged profile around TSS. (**D**) Fold change of H3K27Ac and H3K4me3 ChIP-seq counts within 1kb around TSS for pH-insensitive and pH-antagonistic groups. Welch t-test and Holm-Sidak’s multiple comparisons test, ns p>0.05, * p<0.05, ***p<0.001. (**E**) Profile of ATAC-seq, H3K27 acetylation, H3K4 tri-methylation, p65 and IRF3 ChIP-seq at selected LPS induced genes. Orange boxes highlight regions with strong differences between pH 7.4 and pH 6.5. (**F**) model illustration of the activation of pH-sensitive and pH-insensitive inflammatory response.

A second possibility of a delayed activation kinetics involves chromatin remodeling. Given that acidic pH has been linked to modulation of histone acetylation and metabolism in cell lines,^56,57^ we performed ATAC-seq and ChIP-seq to examine chromatin accessibility and histone modifications (H3K27ac and H3K4me3) associated with transcriptional activation. We first identified ATAC-seq peaks that were significantly induced by LPS (> 4-fold, pH 7.4 LPS vs pH 7.4) or homeostatically maintained (< 2-fold) at pH 7.4 across two biological replicates. At the LPS-induced peaks, we observed a comparable increase in ATAC-seq, H3K27ac and H3K4me3 after LPS between pH 7.4 and 6.5, while the homeostatic peaks remain unchanged among all examined conditions (Figure S3E). Intriguingly, ATAC-seq signals were mildly elevated at pH 6.5 independent of LPS stimulation (Figure S3E), suggesting that acidic pH does not restrict chromatin accessibility. The consistent levels of global H3K27ac and K3K4me3 signals were also aligned with gene-specific control observed at pH 6.5. Next, we analyzed chromatin changes specific to pH^IN^, pH^ANTI^ and pH^SYN^ genes (Figure 3C). Similar to the genomic profile, ATAC-seq signals at TSS were elevated at pH 6.5 across all three gene groups and demonstrated similar increase after LPS between pH 7.4 and 6.5. We also did not observe a consistent difference in H3K27ac. Only the increase of H3K4me3 was selectively reduced in pH^ANTI^ genes after LPS stimulation at pH 6.5, particularly within the gene body immediately downstream of TSS (Figure 3C, D), consistent with reduced transcriptional activity. Intriguingly, the pH^ANTI^ genes exhibited lower H3K27ac and H3K4me3 signals at baseline, albeit similar levels of expression and chromatin accessibility (Figure S3F). Overall, both the lack of pH-dependent difference in global histone modifications and the comparable regulation of chromatin accessibility at TSS suggest that acidic pH may impact distal regulatory elements to control gene activation.

The association between specific enhancers and genes is challenging to define at genome-wide in macrophages. We thus turned to investigate examples of pH-dependent and independent genes. At *Il6* and *Edn1* loci, two representative pH^Anti^ genes, transcriptional induction was correlated with the activation of multiple distal enhancers spanning hundreds of thousands of base pairs—marked by increased ATAC-seq and broadened H3K27ac peaks at pH 7.4 following 4 hours of LPS stimulation (Figure 3E). Notably, the H3K27ac signals at these enhancers were completely abolished at pH 6.5, along with a reduction in chromatin accessibility and binding of p65 and IRF3 by ChIP-seq. In contrast, the pH^IN^ gene *Nfkbia* displayed consistent epigenetic marks across pH conditions, and the pH^SYN^ gene *Ifnb1* exhibited elevated H3K27ac and H3K4me3 at gene TSS, as well as enhanced IRF3 binding at a-15kb enhancer (Figure 3E), arguing against the possibility that acidic pH pleiotropically blocks histone modification or epigenetic remodeling. To probe this further, we inhibited histone acetyltransferase (HAT) p300 with C646 and histone deacetylases (HDACs) with pan-inhibitor TSA.^58,59^ Overall, HAT inhibition resulted in a less profound effect than HDAC inhibition, yet both differed from the influence of acidic pH (Figure S3G). For instance, TSA strongly inhibited both pH^IN^ and pH^SYN^ genes (Figure S3D, S3G), and the activation of pH^ANTI^ genes required both HAT and HDAC activity (Figure S3G). Thus, combining epigenetic profiling and pharmacological perturbations, our data suggest that a pH-sensitive epigenetic mechanism specifically impacts the activation of inflammatory genes. Genes induced immediately by TLR signaling are insensitive to pH difference, while genes dependent on the activation of NF-ΚB, IRF3 and distal enhancers are more likely to be influenced by the tissue microenvironment. A putative transcriptionally induced repressor belonging to the pH^ANTI^ group may be necessary to deactivate *Ifnb1*, *Adm* and other pH^SYN^ genes at pH 7.4; its absence at pH 6.5 leads to their prolonged and heightened activation (Figure 3F).

### Transcriptional condensates of BRD4 are sensitive to acidic pH

The budding yeast Snf5, a core component of SWI/SNF chromatin remodeling complex, was recently found to be sensitive to intracellular pH (pHi).^60^ Two histidine residues on a disordered loop of *S. cerevisiae* Snf5 become protonated at pH 6.5 and this protonation disrupts the electrostatic interactions with nucleosomes and transcription factors.^60^ However, this disordered loop is only present in fungi. Guided by its biochemical properties, we performed a bioinformatic screening to identify putative pH sensors that could mediate pH-dependent transcriptional response in macrophages. Our screening was based on three assumptions: 1) a pH-sensitive protein must carry a peptide region with a significant difference in protonation between pH 7.4 and 6.5 (Δcharge), 2) this region is near non-polar residues or disordered sequences, with minimal prior structural constraints, and 3) this region is enriched in prolines (P) and glutamines (Q), similar to the disordered loop in yeast SNF5. We first scanned 50,961 annotated proteins in the mouse genome (UniProt) to identify putative regions with significant Δcharge (> 1 histidine residues in a stretch of 20 amino acids). We then filtered them based on IDR consensus score^61^ and PQ-enrichment, and gene expression in BMDMs (Figure 4A, Supplemental table 3). BRD4, a BET family member known for recognizing acetylated histones and regulating transcriptional programs in development and inflammation,^46^ was identified as a top candidate. BRD4 has two regions enriched for histidine, proline, and glutamine (HPQ) (Figure 4A): AA 721-800, featuring a stretch of six consecutive histidine residues adjacent to a 40-residue non-polar region with 90% poly-PQ, and AA 1001-1080, containing nine histidine and 39 proline or glutamine residues. These regions have a minimal net charge (Figure S4A) and are within the C-terminal IDR of BRD4 (IDR^BRD4^). The IDR-containing full length BRD4 is the most abundant isoform expressed in BMDMs (Figure S4B, C).

**Figure 4.**
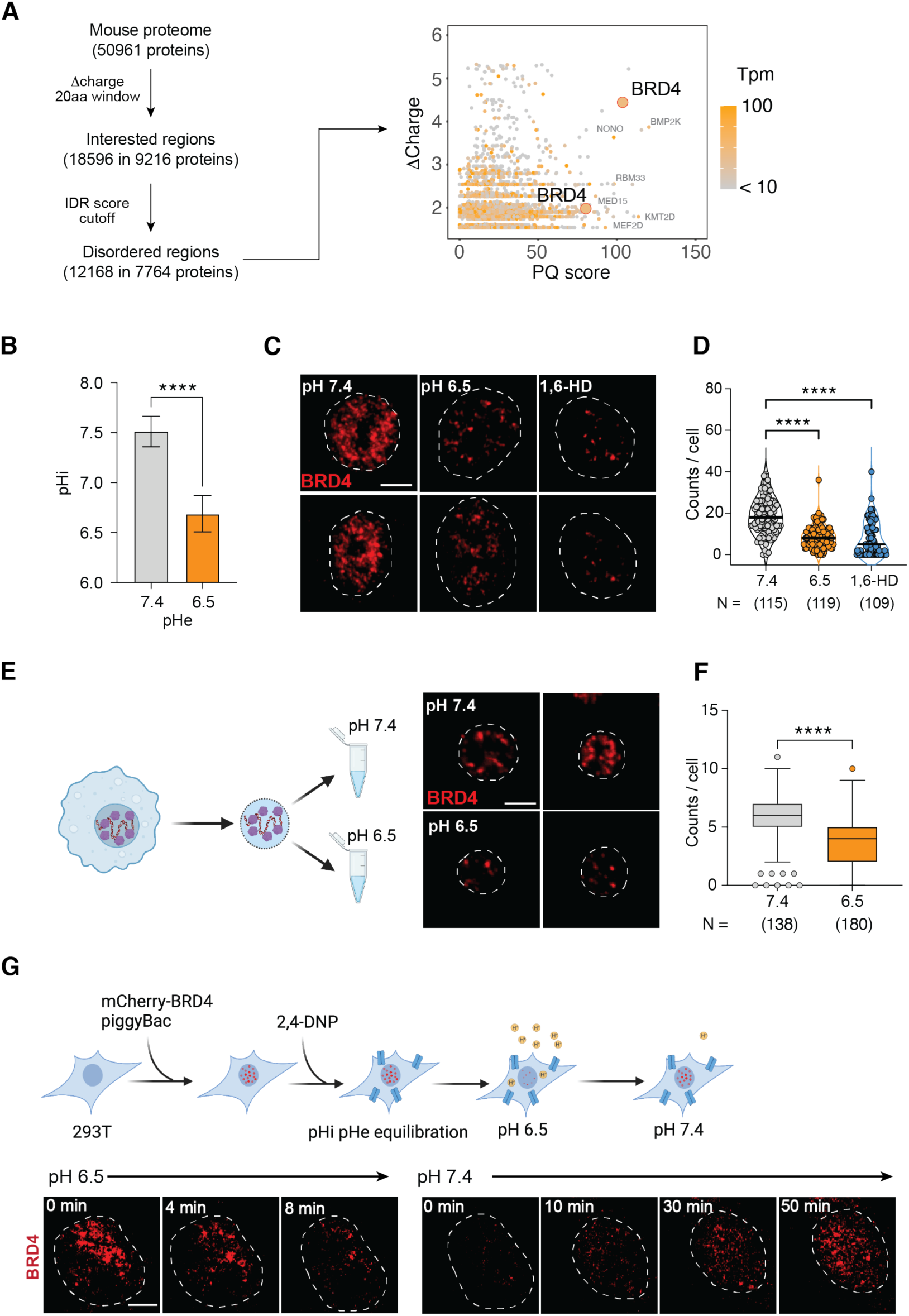
Acidic pH regulates BRD4 transcriptional condensates. (**A**) Diagram and results of the bioinformatic analysis of pH-sensitive peptide sequences in the mouse proteome. The 2D-plot shows side chain Δcharge and enrichment of proline and glutamine residues. Orange color indicates gene expression in BMDMs. (**B**) pHi measurement in BMDMs using pH-sensitive fluorescent probe SNARF-4F. **** p<0.0001, unpaired Student t test. (**C-D**) Immunofluorescent imaging (C) and quantification (D) of BRD4 in BMDMs cultured at pH 7.4, 6.5 for 4 hours, or treated with 10% 1,6-Hexanediol for 2 min. **** p<0.0001, One way ANOVA test. (**E-F**) Immunofluorescent imaging (E) and quantification (F) of BRD4 in isolated BMDM nuclei, conditioned in pH 7.4 and pH 6.5 buffer. **** p<0.0001, unpaired Student’s t test. (**G**) Time lapse imaging of BRD4 in live BMDMs in response to pH 6.5. N represents the number of cells analyzed for each condition in (E, G). Scale bar represents 2 μm.

Using a pH-sensitive fluorescent dye SNARF,^62^ we found that the intracellular pH (pHi) of BMDMs decreased to pH 6.6 in pH 6.5 medium (Figure 4B, S4D), exposing BRD4 to an acidic intracellular environment. Since BRD4 forms transcriptional condensates via its hydrophobic IDR,^33^ we hypothesized that pH-dependent protonation of histidine residues may disrupt the hydrophobic interactions critical for condensate formation. At pH 7.4, endogenous BRD4 formed small and distinct foci in BMDMs, and these foci were substantially reduced at pH 6.5 (Figure 4C, D). Treatment with 10% 1,6-hexanediol (1,6-HD), a chemical known to disrupt weak hydrophobic macromolecular interactions, reduced BRD4 foci similarly to acidic pH. Both BRD4 mRNA and protein expression remained stable at pH 6.5 (Figure S4E, F), demonstrating that condensates rather than BRD4 expression were regulated by pH. To discern potential roles of signaling and cytoplasmic contents, we isolated BMDM nuclei after gently lysing the plasma membrane and incubated them at pH 7.4 or 6.5 for 0.5 hours, followed by fixation and imaging (Figure S4G). Endogenous BRD4 could form distinct foci in isolated nuclei, although fewer in quantities than in living cells (Figure 4F). Despite the morphological difference, we found that BRD4 foci significantly decreased in acidic pH (Figure 4G), demonstrating nuclear intrinsic roles in regulating pH-dependent BRD4 condensates.

To further investigate the dynamics of BRD4 condensates, we generated a stable 293T cell line expressing a murine mCherry-BRD4 fusion protein for live-cell imaging. Initially, we noticed a heterogeneous response in mCherry-BRD4, likely due to variability in pH buffering capacity between cells. Treating 293T cells with a proton ionophore 2-4-Dinitrophenol (2,4-DNP) equilibrated extracellular and intracellular pH,^63^ and significantly reduced the heterogeneity in BRD4 condensates (Figure S4H). To monitor the dynamic changes, we treated these cells with 2,4-DNP for 30 min at pH 7.4 and abruptly switched the medium to pH 6.5. Remarkably, BRD4 condensates were significantly reduced within 10-12 minutes, and upon returning to pH 7.4, the condensates reappeared and recovered in 1 hour (Figure 4H, S4I, Supplemental movie 1). In contrast, cells expressing only mCherry showed constant fluorescent signals unaffected by pH (Figure S4J).^64^ Collectively, these findings demonstrated that BRD4 condensates are dynamically regulated in a pH-dependent manner within live cells, highlighting their potential role in regulating cellular responses to environmental shifts.

### Interference with BRD4 functions largely recapitulates pH-dependent responses

Given that pH-dependent changes in BRD4 condensates are reversible, we investigated whether the impact on inflammatory responses can be reversed after normalizing pH. BRD4 condensates were significantly reduced after overnight incubation at pH 6.5, and were fully restored after 4 hours at pH 7.4 (Fig. 5A, B). Alongside the changes in BRD4 condensates, pH-repressed genes regained activation nearly completely after re-conditioning at pH 7.4 (Fig. 5C). To test whether pH-dependent genes are regulated by BRD4, we analyzed the transcriptional response to LPS in *Brd4*^-/-^ BMDMs.^65^ We applied the deconvolution model to identify LPS-induced genes that are BRD4-dependent and independent, and found that the BRD4-dependent genes were more repressed under acidic conditions (Figure 5D). Similarly, competitive inhibitors targeting the bromodomains of BRD4 (JQ-1, iBET, and MS-645), or disrupting condensates with 1% 1,6-HD, consistently repressed pH^ANTI^ genes (Figure 5E). Thus, the pH-dependent inflammatory regulation is functionally linked to BRD4 condensates.

**Figure 5.**
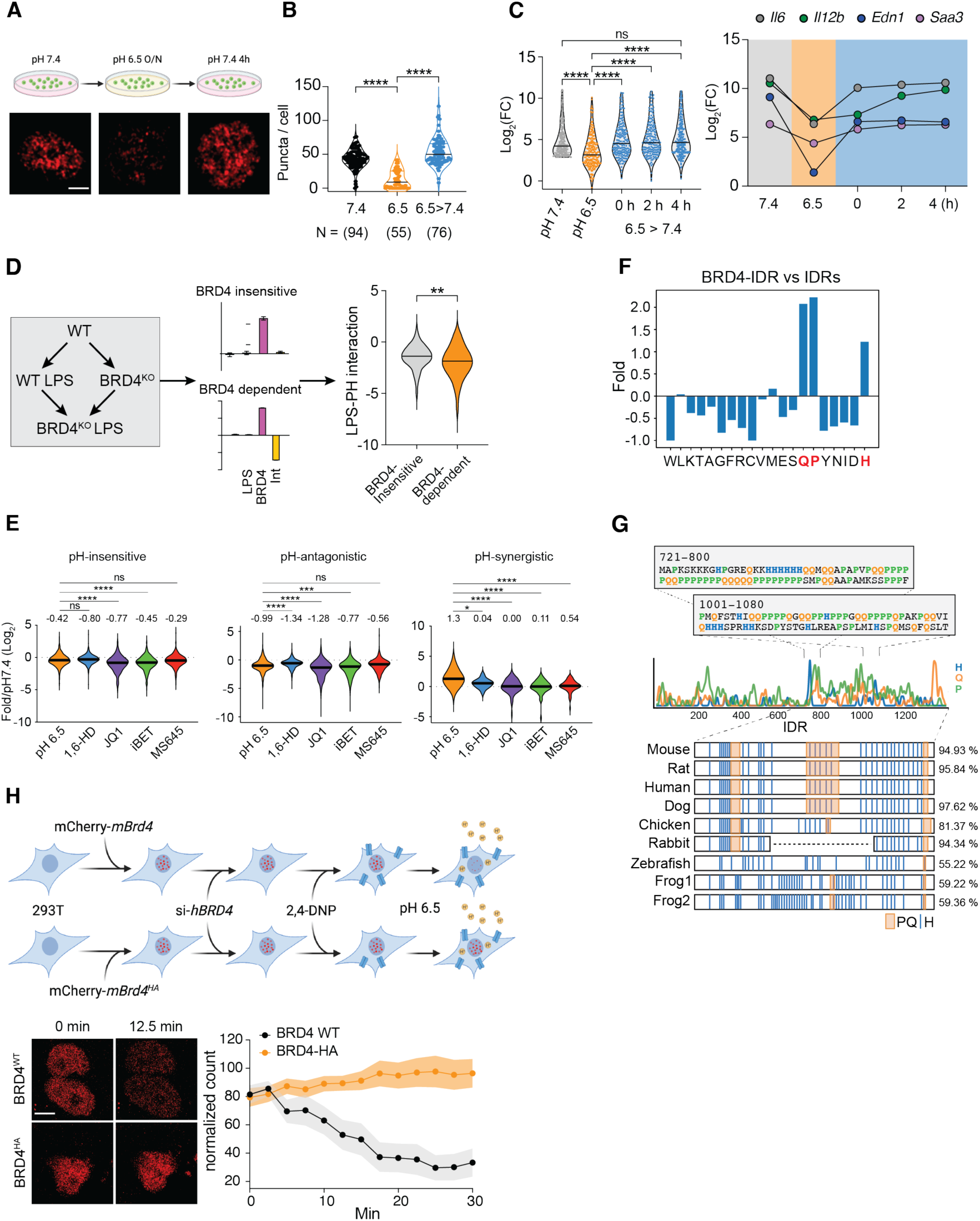
BRD4 is a pH sensor and regulates pH-dependent inflammatory response. (**A-B**) Immunofluorescent imaging (A) and quantification (B) of BRD4 in BMDMs. Brown-Forsythe and Welch ANOVA tests. (**C**) Violin plots of fold activation for pH-sensitive genes after restoring media pH from 6.5 to 7.4. The top 200 pH-repressed genes. Median of fold activation is labeled on violin plots. Kruskal-Wallis test with multiple comparisons. Only significant pairs are labeled. (**D**) RNA-seq deconvolution analysis of BRD4 KO BMDMs stimulated with 10 ng/mL LPS for 4 hours. LPS-induced BRD4-dependent and independent genes were analyzed for their pH-dependence. Unpaired Student’s t test. (**E**) Fold difference of pH-insensitive,-antagonistic or synergistic genes, at pH 6.5, with 1% 1,6-hexanediol or various BRD4 inhibitors. (**F)** Amino acid composition in the BRD4-IDR relative to that of all annotated IDRs in the mouse proteome. (**G**) Conservation of HPQ regions along the coding sequence of BRD4. Panels display amino acid sequences of the two HPQ regions within BRD4-IDR identified through bioinformatic screening. The distribution of H, P, Q residues, along with HPQ patterns in BRD4-IDR across different vertebrates are illustrated with sequence conservation. (**G**) Live-cell imaging of mCherry-BRD4^WT^ or mCherry-BRD4^HA^ 293T cells in response to pH 6.5. N represents the number of cells analyzed for each condition (B, C, D, E). ns, p>0.05, *p<0.05, **p<0.01, ***p<0.001, **** p<0.0001.

Histidine is responsible for the majority of protonation around physiological pH. We found that IDR^BRD4^ is uniquely enriched for histidine (Figure 5F, S5A), as both full length BRD4 (Figure S5B) and IDRs from mouse proteome (Figure S5C) lack histidine enrichment. Moreover, the two identified HPQ regions are highly conserved among vertebrates (Figure 5G), suggesting the biological significance of these sequence features. To investigate whether protonation at histidine residues directly contribute to pH-dependent condensates, we synthesized a BRD4^HA^ mutant that swaps 33 histidine residues within IDR^BRD4^ to alanine, to eliminate most pH-dependent Δcharge (Figure S5D). In 293T cell lines expressing comparable levels of murine mCherry-BRD4^WT^ or mCherry-BRD4^HA^ (Figure S5D), we observed that both BRD4 variants appeared in condensates, likely due to high sequence homology between mouse and human. To minimize the complication of endogenous human BRD4, we performed all live-cell imaging at 24 hours after siRNA knock-down of hBRD4. In these experiments, we observed that the number of BRD4^WT^ condensates (36 cells) began to decrease between 2-5 minutes at pH 6.5 (Supplemental movie 2), stabilizing at ∼25% of their original quantity by half an hour; in contrast, these pH-dependent changes were completely abolished for BRD4^HA^ (45 cells) (Figure 5H, Supplemental movie 3). These data strongly support that BRD4 directly senses intracellular pH via its histidine-enriched IDR, thus dynamically regulating transcriptional condensates to control gene expression.

### Gene-specific mechanisms underlie pH-dependent regulation

BRD4 binds to and regulates both pH-sensitive and pH-insensitive genes (Figure 5E). Then how could pH-dependent BRD4 condensates specifically influence a subset of inflammatory genes? The mainstream model of BRD4-dependent gene activation illustrates that BRD4 recognizes histone acetylation via its bromodomains and recruits mediator complexes to active enhancers, facilitating the interaction between distal enhancers and promoters to activate transcriptional machinery.^39^ Given the distinct changes at enhancers of pH-sensitive genes, we hypothesize that pH-dependent condensates facilitate remodeling and activation of enhancers, and thus control gene activation that strongly depends on these processes.

First, bridging enhancer activation to gene promoters requires mediators.^66^ Both BRD4 and MED1 form liquid-liquid phase condensates, and co-localize at distinct nuclear foci to recruit RNA polymerase II.^33^ At pH 7.4, endogenous MED1 formed condensates and co-localized with BRD4 in BMDMs (Figure 6A). At pH 6.5, however, these condensates were substantially reduced, despite MED1 lacking an HPQ-enriched IDR (Figure S6A, B). Notably, the remaining puncta of BRD4 and MED1 were partitioned spatially, contrasting with the strong colocalization seen at pH 7.4 (Figure 6A, B). These pH-dependent changes were also aligned with chemical disruption of condensates by 1,6-HD (Fig. 6A, B). Since reduction of H3K27Ac was observed at the enhancers of pH-sensitive genes, we thus tested whether interfering binding to histone acetylation would result in a similar perturbation of BRD4 condensates. Surprisingly, BRD4 and MED1 condensates were differentially affected by the bromodomain competitive inhibitors JQ1, iBET and MS645, although all of them inhibited BRD4 functions (Figure 5E). The contrasting phenotypes between acidic pH and BRD4 inhibitors also implied that altered chromatin recruitment was not the cause for pH-dependent dissolution of transcription condensates. This is corroborated by comparable levels of global histone acetylation (Figure S3E), binding of BRD4 at house-keeping genes (Figure S6C), and previous findings that the reliance of condensates on histone acetylation may vary by cell type.^33,41,67^ Thus, acidic pH disrupts the formation of transcriptional condensates containing both BRD4 and MED1, two essential components of enhancer-regulated transcriptional activation.

**Figure 6.**
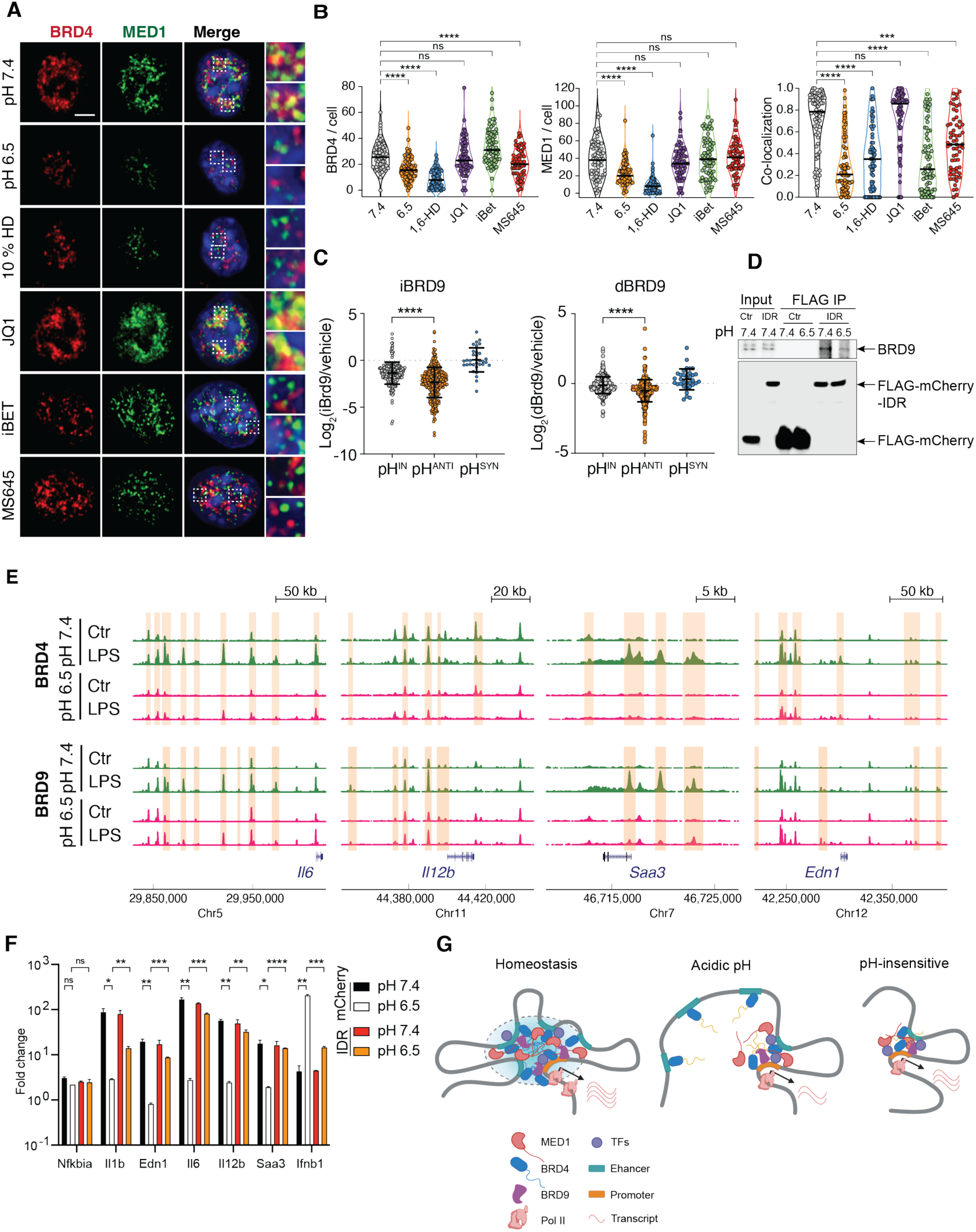
Gene-specific regulation is mediated by BRD4-MED1 transcription condensates and BRD9. (A-B) BRD4 and MED1 in BMDMs at pH 7.4, at 6.5, in the presence of 10 % 1,6-HD, JQ-1, iBET and MS645. (A) Immunofluorescent imaging (B) Quantification of puncta and co-localization. One way ANOVA. (**C**) Fold change of LPS induced gene expression in the presence of BRD9 specific inhibitor (iBRD9) of degrader (dBRD9). (**D**) Western blot analysis of Co-IP between BRD4-IDR and BRD9. FLAG-mCherry was fused with BRD4-IDR for IP with total cell lysate, with FLAG-mCherry as control (Ctrl). (**E**) ChIP-seq of BRD4 and BRD9 in BMDMs at pH-regulated genes. Orange boxes highlight regions with significant differences in BRD4 or BRD9 binding. (**F**) Fold change of pH-dependent inflammatory genes in iBMDMs overexpressing BRD4-IDR after 6 hours 100 ng/mL LPS at pH 7.4 or 6.5, normalized to unstimulated conditions respectively. Mean+/-standard deviation (STD). Unpaired t-test, Holm-Sidak’s test for multiple comparisons. (**G**) Model diagram of pH-dependent regulation of BRD4 condensates. In (B), (D), (F) ns p>0.05, *** p<0.001, **** p<0.0001.

Second, enhancer activation often requires chromatin remodeling. Bromodomain-containing protein 9 (BRD9), a subunit of the noncanonical SWI/SNF chromatin remodeling complex (ncBAF), was found to co-localize with BRD4 at the enhancers and promoters enriched with NF-ΚB and IRF3 binding motifs.^68^ The inhibition of BRD4 with JQ1 displaces BRD9 in both BMDMs and embryonic stem cells ^68,69^ without directly inhibiting BRD9 *in vitro*.^70^ These findings suggest that BRD4 can recruit BRD9-containing ncBAF complex to activate enhancers. We hypothesized that disruption of BRD4 condensates leads to a defective recruitment of BRD9 to regulate pH-sensitive genes. Indeed, in the presence of a specific BRD9 inhibitor (BRD9i) or a BRD9 degrader (dBRD9), pH^ANTI^ genes exhibited a significantly reduced activation in comparison to pH^IN^ genes (Figure 6C).^68^ ChIP-seq analysis of BRD4 and BRD9 revealed a significant reduction in LPS-induced recruitment at the enhancers and promoters at pH 6.5, particularly for pH-sensitive genes such as *Il6, Il12b, Saa3, Edn1* (Figure 6E). Among BET family proteins, only BRD4 contains HPQ regions, suggesting that pH-dependent BRD9 recruitment is likely mediated through BRD4 (Fig. S6A, B). A recent proteomic analysis further suggested that the IDR^BRD4^ may directly interact with BRD9.^71^ We thus expressed FLAG-mCherry-IDR^BRD4^ to test whether the recruitment BRD9 is affected by pH. We observed that FLAG-mCherry-IDR^BRD4^ were integrated into endogenous BRD4 condensates (Figure S6D). Using Co-immunoprecipitation (Co-IP), we found that the co-association of BRD9 to BRD4 was substantially reduced at pH 6.5 (Figure 6D). In connection with the differential regulation of distal enhancers of pH-sensitive genes (Figure 3E), these data suggest that pH-dependent BRD4 condensates may regulate a subset of inflammatory response in macrophages by directly recruiting the non-classic BAF complex to activate enhancer.

BRD4, MED1 and enhancer activation are crucial for RNA Pol II elongation. We speculate that Pol II elongation at pH-sensitive genes was selectively inhibited at acidic pH conditions, and activating elongation may mitigate the suppressive inflammatory response. The C-terminal fragment of BRD4 was found sufficient to release paused Pol II independent of the BET bromodomains.^72^ We thus tested the possibility of rescuing pH-dependent repression using IDR^BRD4^. In an immortalized BMDM cell line (iBMDMs), we identified pH sensitive genes consistently regulated with BMDMs (Figure S6E). Over-expression of FLAG-mCherry-IDR^BRD4^ significantly reversed the pH-dependent repression and reduced the synergistic induction of *Ifnb1*, while maintaining proper activation of pH-insensitive genes (Figure 6F). Thus, we concluded that transcription condensates integrate environmental signals to orchestrate a gene-specific regulation of inflammatory response facilitated by chromatin remodeling (Figure 6G). Leveraging BRD4-IDR can reverse environment-dependent repression of inflammatory responses.

### pH sensing by BRD4 mediates feedback control of inflammatory activation

To understand the potential functions of pH-dependent transcriptional condensates during inflammation, we revisited the change of pH upon stimulation with LPS. Interestingly, we found that the pHi of BMDMs quickly decreased and plateaued after 8 hours (S7A), in addition to acidifying the extracellular environment (Figure 1B). The decrease in pHi cannot be simply restored by conditioning LPS-stimulated BMDMs in pH 7.4 medium (Figure 7A), suggesting that LPS activates an intrinsic program to maintain a different intracellular state. Consequently, BRD4 and MED1 condensates were significantly reduced simply by the activation of innate sensing pathways (Figure 7B and C). We found that subjecting LPS-treated BMDMs to a 5 min pulse of 10 μM nigericin and 100 mM KCl followed by conditioning at pH 7.4 can largely restore the pHi to 7.0 (Figure 7A). Following this treatment, the increase in pHi restored both BRD4 and MED1 condensates as well as their colocalization (Figure 7B and C), demonstrating that low pHi induced by LPS *in vitro* is sufficient and necessary to inhibit transcriptional condensates. *In vivo*, we examined the response of thioglycolate-induced peritoneal macrophages (pMac) to i.p. Injection of LPS. Thioglycolate induces the expansion of CD11b^Int^ F4/80^low^ MHC-II^+^ pMac (Figure S7B).^73^ Although the number of pMac decreased after LPS treatment, the remaining pMac from peritoneal cavity displayed a significant reduction in pHi and BRD4 condensates, compared to the PBS-treated controls (Figure 7D, E). Altogether, these data demonstrated that inflammatory activation of macrophages *in vitro* and *in vivo* adopts an acidic intracellular environment that is necessary and sufficient to regulate transcription condensates. Since acidic pH represses the inflammatory response in macrophages, our data suggest that sensing pH via BRD4-containing transcription condensates function as negative feedback to control the inflammatory response.

**Figure 7.**
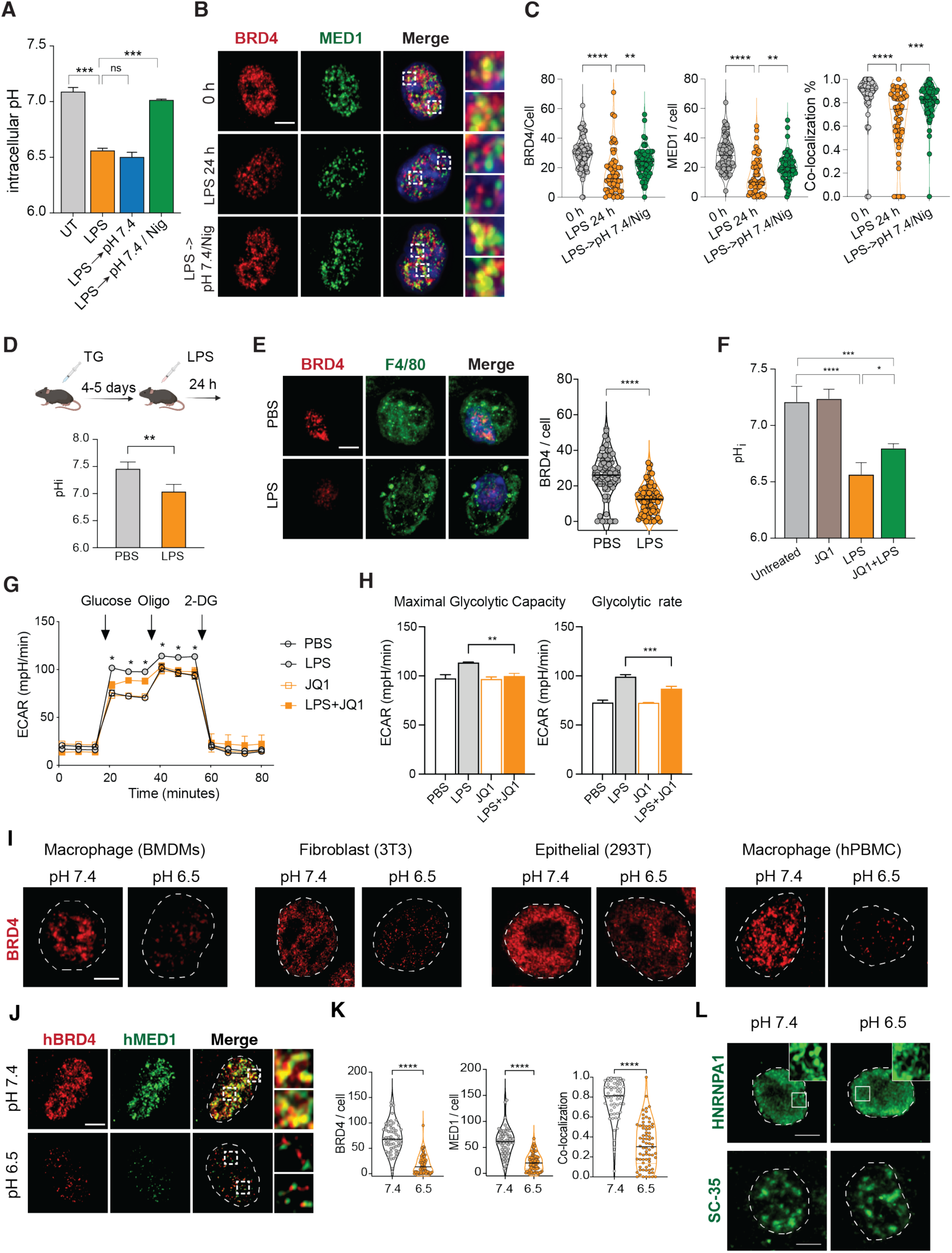
pHi sensing via BRD4 acts as negative feedback to modulate pH homeostasis and inflammatory response. (**A)** pHi in BMDMs after LPS stimulation in various indicated conditions, pairwise comparison with Kruskal-Wallis. (**B-C**) Immunofluorescent imaging (B) and quantification (C) of BRD4, MED1 and co-localization. one way ANOVA (F). (**D-E**) pHi and BRD4 measurement of thioglycollate-induced peritoneal macrophages *in vivo*, 24 h after i.p. injection with PBS or 3 mg/kg LPS. pH quantification, unpaired Student’s t test (D), Immunofluorescent imaging and quantification of F4/80+ macrophages, Mann-Whtiney test (E). (**F**) pHi of BMDMs treated with 100 ng/mL LPS for 24 h and/or 0.5 μM JQ1 for 8 h, one way ANOVA. (**G**) Seahorse analysis of glycolytic activity in BMDMs treated with 100 ng/mL LPS and/or 0.5 μM JQ1 for 8 h, one way ANOVA. (**H**) Glycolytic capacity and rate of BMDMs treated with 100 ng/mL LPS +/-0.5 μM JQ1 for 8 h, one way ANOVA. (**I, L**) Immunofluorescent imaging of BRD4 (I), HNRNPA1 or SRSF2 (L) phase condensates in murine and human cells. (**J, K**) BRD4 and MED1 puncta in human macrophages at pH 7.4 and pH 6.5. (J) Immunofluorescent imaging, (K) Quantification of puncta and co-localization. Mann-Whtiney test. p>0.05, *p<0.05, **p<0.01, ***p<0.001, **** p<0.0001.

To explore the role of pH-sensing by BRD4 in cellular physiology, we reasoned that a pH sensor may regulate cellular processes to control pHi, similar to a homeostatic controller (Figure S7C).^1,3^ Thus, we tested whether BRD4 regulates LPS-induced intracellular acidification. Interestingly, JQ-1 treatment alleviated LPS-induced acidification in BMDMs by 0.23 pH unit (1.44-fold of protons) without impacting the pHi in naive BMDMs (Figure 7F). Using seahorse assay, we observed that the LPS-induced increase in both glycolytic capacity and glycolytic rate was impaired by JQ-1 (Figure 7G, H), similar to BMDMs activated at pH 6.5 (Figure S7D). Mechanistically, we found that hexokinases (HK1, HK2, HK3) that catalyze the phosphorylation from glucose to phopho-6-glucose, the first chemical reaction of converting glucose to energy, were transcriptionally induced by LPS in BMDMs in a BRD4-dependent manner (Figure S7E). Thus, BRD4 acts as a sensor for pHi to modulate both transcriptional and metabolic inflammatory response.

### BRD4 is a generic pH sensor in multiple cell types

Finally, given the broad expression and crucial role of BRD4 in many cell types, we investigated if pH-dependent condensates is a general phenomenon across various cell types and species. We observed that murine and human primary macrophages, as well as stromal and epithelial cell lines, all exhibited pH-dependent regulation of BRD4 condensates (Figure 7I). In particular, BRD4 and MED1 in human macrophages derived from peripheral blood mononuclear cells displayed extremely strong pH dependence (Figure 7J, K). It is worth noting that different cell types may differ in their sensitivity to extracellular pH and likely exhibit cell-type dependent response under acidic environment *in vivo*. Moreover, we found that other types of nuclear condensates, such as RNA granules and stress granules were not sensitive to the pH conditions examined here (Figure 7L). Thus, pH-sensing by BRD4 condensates establishes a unique mechanism to integrate extracellular and intracellular environments in regulating inflammatory response and beyond.

## Discussion

During inflammation, extreme perturbations in cellular or tissue microenvironment signals strong deviation from tissue homeostasis. Adjusting immune responses to the changes in the microenvironment may provide an adaptable strategy for the immune system to calibrate inflammatory responses to the consequences of inflammation.^2,3^ Inflammatory response leads to acidification of both tissue and cellular microenvironment. In this work, we found that tissue pH regulates a switch-like transcriptional response in macrophages. Mechanistically, acidic pH disrupts transcription condensates of BRD4 and MED1 that are crucial for activating a sub-set of inflammatory genes requiring enhancer-dependent transcriptional activation. The acidification triggered by microbial cues is both necessary and sufficient to disrupt BRD4 condensates *in vitro* and *in vivo*. Thus, pH-dependent control of transcriptional condensates establishes a new mode of inflammatory regulation, allowing tissue sentinels to directly integrate environmental information to adjust the magnitude and quality of an inflammatory response.

Changes in gene expression in response to external signals are thought to be driven primarily by transcription factors. Thus, macrophage response to LPS is mediated by several transcription factors that cause dramatic changes in gene expression.^21,46^ Upon activation, transcription factors immediately downstream of TLR4 signaling orchestrate waves of sequentially induced autocrine signals along with feedforward or feedback transcriptional regulators, culminating in a dynamic and robust transcriptional response within the first few hours.^50,74^ The LPS-response is comprised of multiple related and coordinated inflammatory functions related to host defense.^48^ However, we found extensive transcriptional reprogramming and switch-like expression patterns in macrophages at different pH levels, without apparent difference in TLR signaling. This observation suggests that innate immunity can be highly adaptable and tunable by the tissue microenvironment. Interestingly, pH-insensitive and pH-sensitive genes appears to partition into functions associated with the kinetics and magnitude of inflammatory response. The early and pH-independent response triggered by LPS primarily involves antimicrobial defense, likely regulated directly by TLR signaling to scale with the inflammatory demand. On the contrary, genes amplifying the inflammatory cascade - such as chemokines, cytokines, antigen presentation molecules and acute phase proteins - are pH-sensitive, executing tasks that may impose a significant negative impact on tissues. The attenuated activation of these genes under inflammatory conditions that significantly induce changes in tissue state, is a strategy that likely evolved to limit the cost of inflammation. Thus, the pH-dependent regulation of inflammatory response in macrophages highlights a potential strategy to decouple the demand of inflammatory response from the cost of inflammation.

Several stress-induced biomolecular condensates are regulated by acidic pH. In the budding yeast, RNA binding proteins Pub1 and Pab1, and prion-like protein Sup35, form condensates at a pH range of 5 to 6, mediated by disordered sequences enriched for negatively charged amino acids (glutamate pKa∼4.25 and aspartic acid∼3.86).^75–77^ A similar mechanism was found in mammalian stress granule protein G3BP1, suggesting a conserved feature to partition macromolecules under extreme conditions.^78^ Other condensate proteins, including 53BP1, FUS and a-synuclein, can transit from a soluble phase to a liquid-liquid phase separation or a condensate state at a similar acidic pH range, but the molecular underpinning of such transition is not well understood.^79,80^ Our work identified that histidine-enriched IDR of BRD4 functions to regulate transcriptional condensates under physiological pH range *in vitro* and *in vivo*. Notably, a motif of consecutive or concentrated histidine residues (H-block) surrounded by a stretch of proline and glutamine residues (PQ-block), likely mediates pH-sensing capacity of IDR^BRD4^. The abundant proline and glutamine residues may establish an unstructured and neutral environment that is necessary for the condensate formation of BRD4 at normal pH, and at the same time, signify any change of the electrostatic interactions created by histidine ionization. At pH 6.5, the positively charged H-block destabilize the multivalent interactions of IDR, leading to a phase transition in transcriptional condensates. Interestingly, H-and PQ-blocks in IDR^BRD4^ present a unique sequence feature contrasting histidine-rich phase-separating proteins. For instance, the formation of filaggrin-containing keratohyalin granules during keratinocyte differentiation is dependent on liquid-liquid phase separation of histidine-rich filaggrin, but in this case, acidic pH promotes droplet formation rather than disrupting it.^81^ In *Drosophila*, A 31-residue peptide (7 Hs, 21 Qs, 0 P) in the neuronal amyloid protein Orb2 forms a beta sheet structure that assembles into stable amyloid fibers as a mechanism of long-term memory.^82^As proline strongly disfavors the formation of a beta sheet, the HPQ region found in IDR^BRD4^ may enable dynamic assembly and disassembly of transcription condensates that are necessary for gene activation, while maintaining environment sensitivity. In fact, H-blocks can be used to engineer synthetic peptides sensitive a pH shift of just 0.3 units.^83^ The sequence feature uncovered here may provide guidance for the discovery of additional pH-sensitive proteins and synthetic design of pH-dependent regulators.

The ability to form molecular condensates have emerged as a unique feature of many transcription and chromatin regulators.^24,25,27^ Recent studies begun to shed light on the regulation of these condensates and their specific roles in gene expression. The IDR of MED1 features alternating blocks of positively and negatively charged residues, which recruit subunits of RNA Pol II and exclude negative transcription regulators to promote transcription.^28^ The chromatin remodeler cBAF subunit ARID1A/B contains blocks of alanine, glycine, and glutamine that are essential to interact with a network of transcriptional regulators.^30^ These findings suggest that transcriptional condensates are heterogenous, partitioned and self-regulated to carry out essential gene expression programs. However, whether such crucial process in cells is regulated by internal or external environment remains unknown. Our current study demonstrates that pH controls the formation of transcriptional condensates in primary human and mouse cells. BRD4 acts as a pH-dependent switch that integrates both extracellular environment and intracellular metabolic state to control the expression of inflammatory genes. These findings establish the environment-dependent transcriptional condensates as a new mode of gene regulation parallel to the classic paradigm driven by receptor signaling, transcriptional factors and chromatin accessibility. It is worth noting that pH-dependent transcriptional response only partially overlaps with BRD4-dependent transcriptional response. Many BRD4-dependent genes are insensitive to acidic pH, suggesting that the transcriptional functions related to transcription condensates per se may be distinct from the functions of condensate components. Moreover, we also observed that inflammatory genes relying on chromatin remodeling at distal enhancers were most sensitive to pH and BRD4 condensates (e.g. *Il6*, *Il12b*, *Edn1*). These genes have multiple enhancers or a long-stretch of enhancer regions specifically after LPS stimulation, mimicking super enhancers found at the lineage-determining genes. These enhancers are likely responsible for establishing inducible microdomains via chromatin loops with gene promoters. Our results support a model that BRD4-containing transcriptional condensates are crucial for remodeling and activating distal enhancers, thereby establishing new enhancer-promoter contacts that facilitate efficient transcriptional elongation. Although our understanding of gene-specific regulation remains incomplete, the power to reverse pH-dependent gene regulation by IDR^BRD4^ suggests that targeting pH-dependent condensate may be a viable strategy to reverse immune suppression associated with acidic tissue microenvironment.

Last, since BRD4 is ubiquitously expressed, pH-dependent regulation of transcriptional condensates may be applicable to mediate cellular responses under acidic environment in various mammalian cell types. In the immune system, such mechanism may impact broadly the specific types of activation, differentiation and polarization programs. In non-immune cells, it may selectively diminish the response to inflammatory cues, potentially allowing cells to tolerate a higher level of inflammatory environment that would otherwise be detrimental.

## Limitations

Significant additional investigation will be needed to define the similarities and differences between these frameworks and the specific impact of pH on inflammatory response triggered by other pattern recognition receptors, in tissue resident macrophages or other cell types, or in the context of different tissues. We observed variable sensitivity to the decrease in extracellular pH among different cell types. The pH-dependent transcriptional control may be cell-type specific within a local tissue microenvironment. Moreover, it remains to be determined how BRD4 transcriptional condensates specifically regulate enhancer activity and/or loop interactions between enhancers and promoters, or how histone modifications and co-condensate molecules impact the sensitivity of pH-dependent regulation. Furthermore, it remains to be investigated whether genetic or synthetic approaches that reverse pH-dependent transcriptional condensate formation present a viable strategy to modulate inflammatory responses in disease settings.

## Acknowledgement

We thank both the current and former Medzhitov lab and the Zhou lab members for discussion on this project. We acknowledge image analysis assistance from the Center for Open Bioimage Analysis (COBA) which is supported by National Institute of General Medical Science NIH P41 GM135019. We thank Y. Peng for generous gift of reagents, Y. Shuang, C. Zhang, C. Annicelli and S. Cronin for supporting the Medzhitov lab animal colony. This work was supported by New Science Initiative to D.L.; T32HL116275 to D.O.; NIH RO1AI167993, R37AI116550, and P30DK34854 to J.C.K.; NIH RO1AI151123, the Pew-Stewart Scholars for Cancer Research and the American Cancer Society Research Scholar Award to D.C.H.; Else Kröner Fresenius Prize for Medical Research, the Blavatnik Family Foundation, the Food Allergy Science Initiative, the Howard Hughes Medical Institute to R.M.; Jane Coffin Childs Memorial funds, NIH R35GM151000 and P30DK034854, the Kenneth Rainin foundation Innovator award, Charles H. Hood Foundation and the G. Harold & Leila Y. Mathers Foundation to X.Z..

## Supplemental movies

Supplemental movie 1: Live-cell time-lapse imaging of mCherry-BRD4 at pH 6.5 and recovery at pH 7.4

Supplemental movie 2: Live-cell time-lase imaging of mCherry-BRD4^WT^ at pH 6.5 Supplemental movie 3: Live-cell time-lase imaging of mCherry-BRD4^HA^ at pH 6.5

## Supplemental tables

Supplemental table 1: qPCR primers used in this study

Supplemental table 2: Gene expression and expression components from deconvolution analysis. The table includes both tpm and read count expression of BMDMs at pH 7.4, pH 7.4 LPS 4h, pH 6.5, pH 6.5 LPS 4h, fold changes, expression components (beta), covariance, p-values and R-square of the linear fitting, and gene cluster number.

Supplemental table 3: Bioinformatic screening of putative pH-sensitive proteins in the mouse proteome.

### STAR★Methods

#### Key resource table

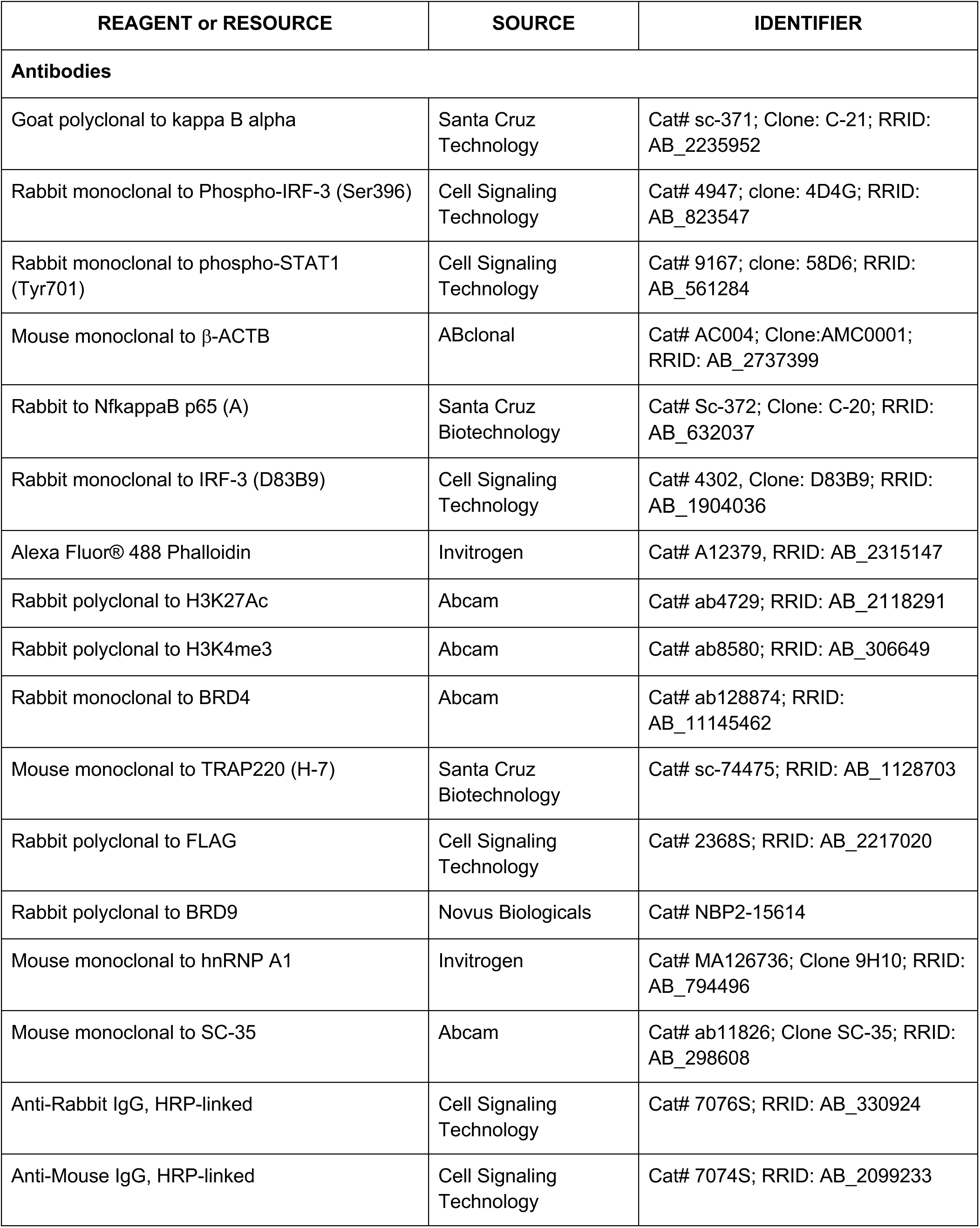
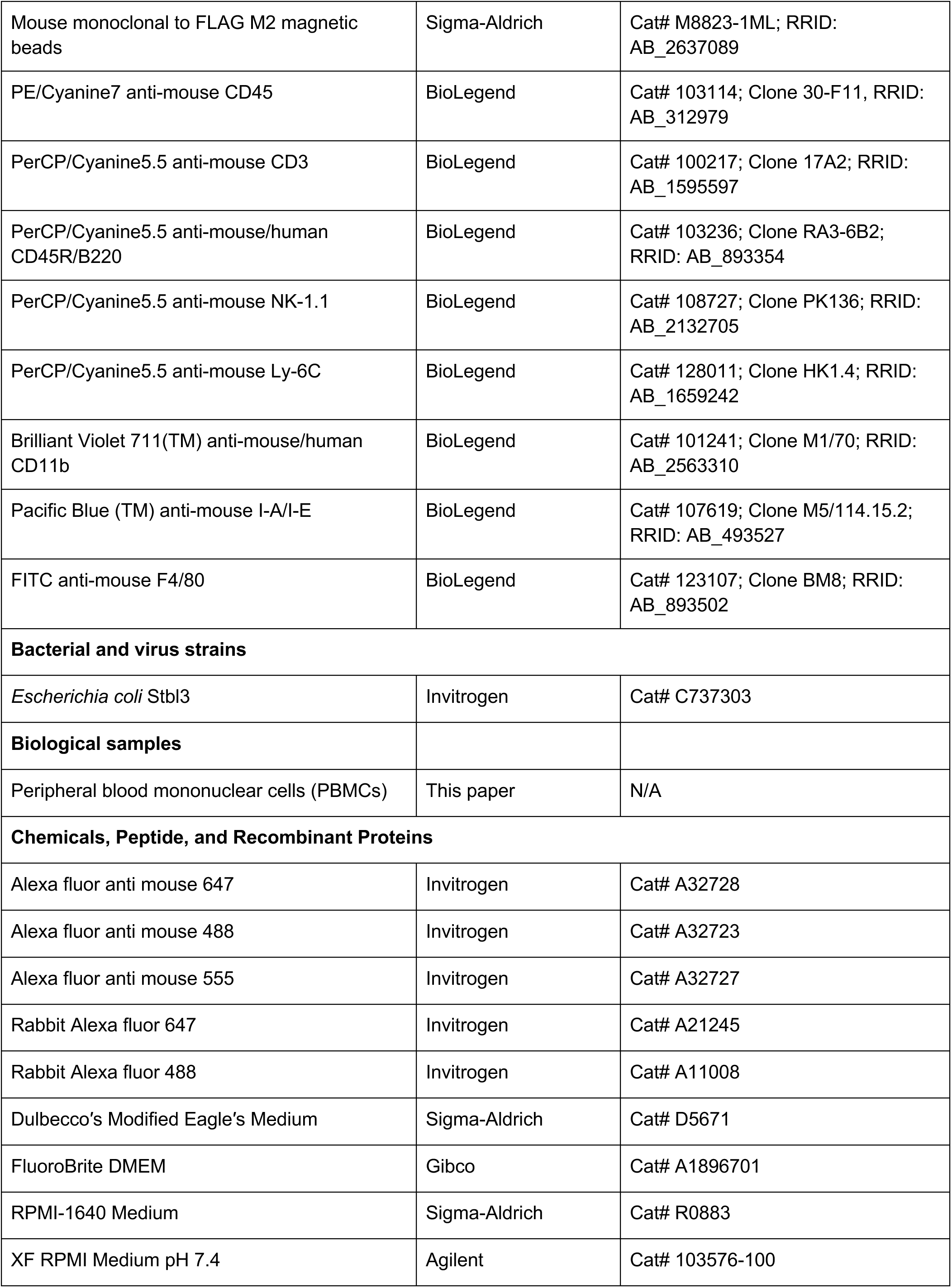
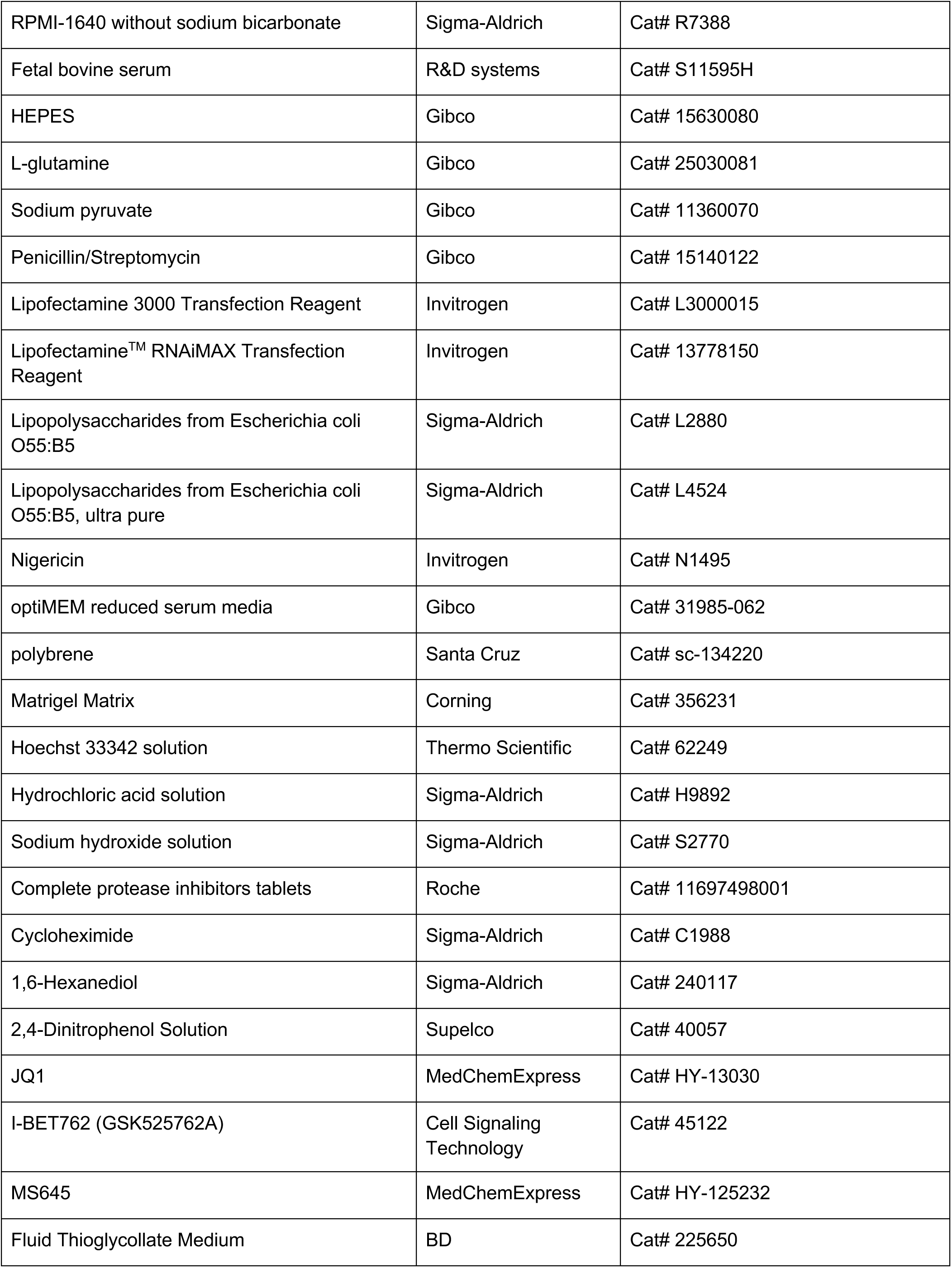
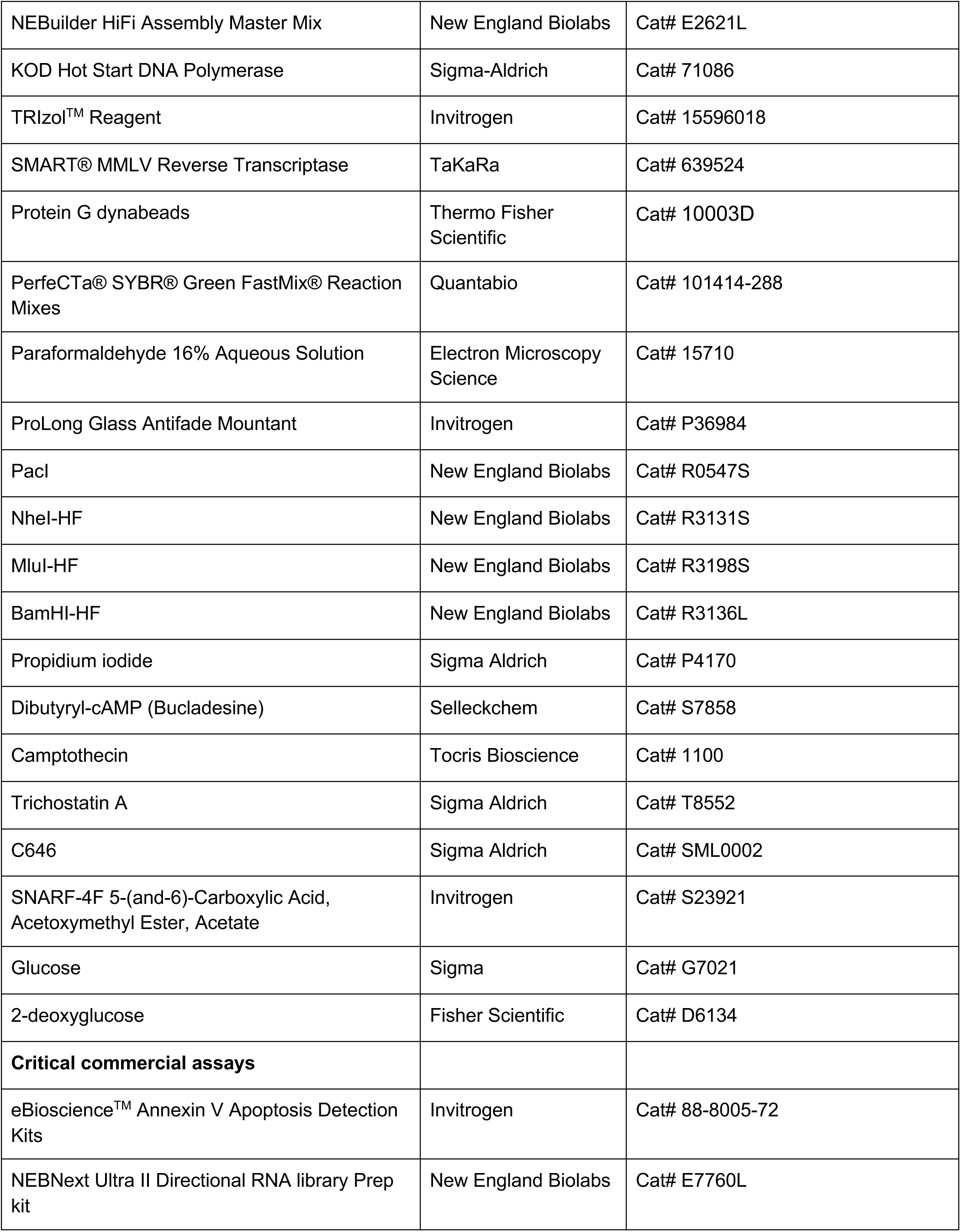
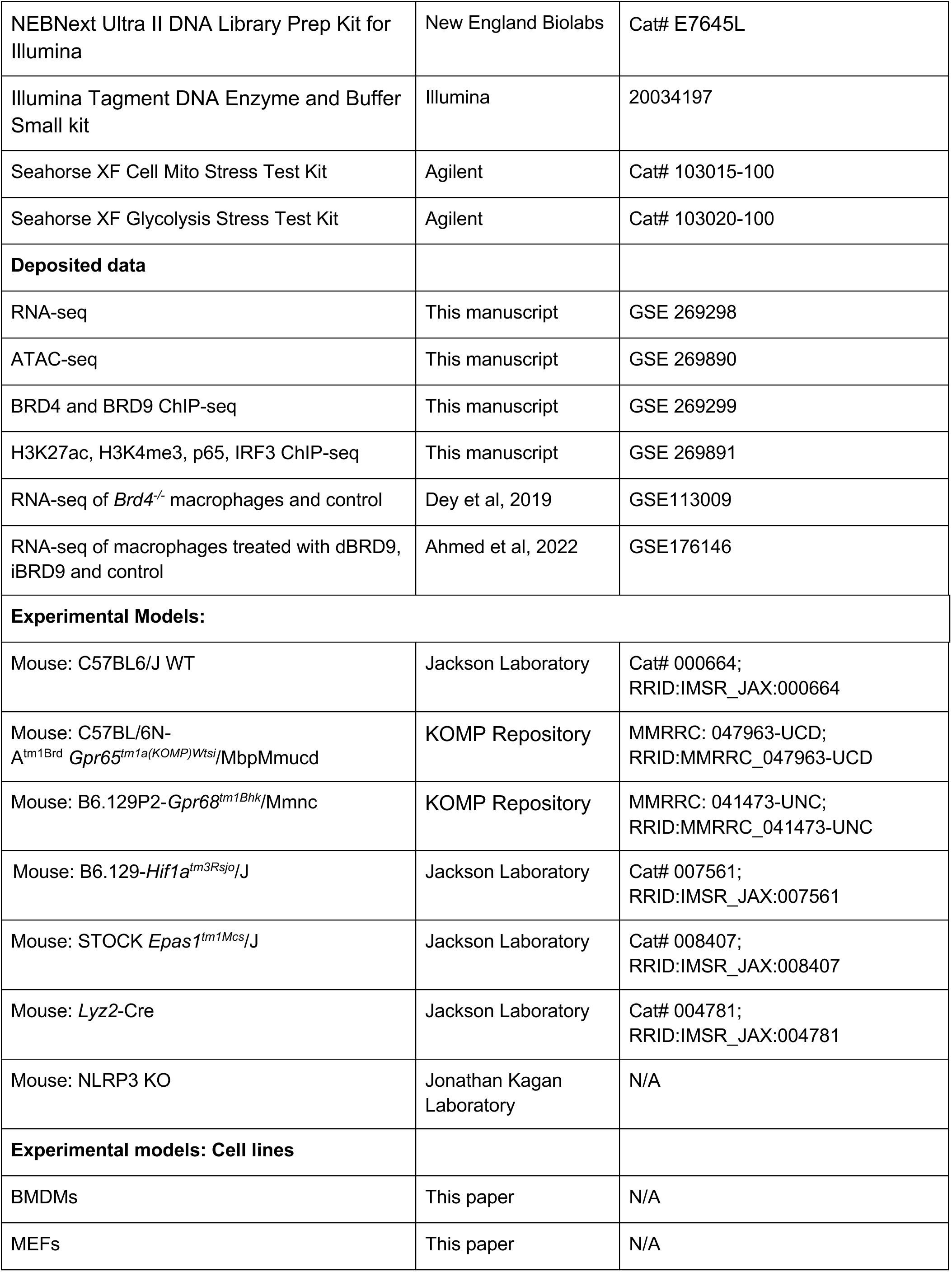
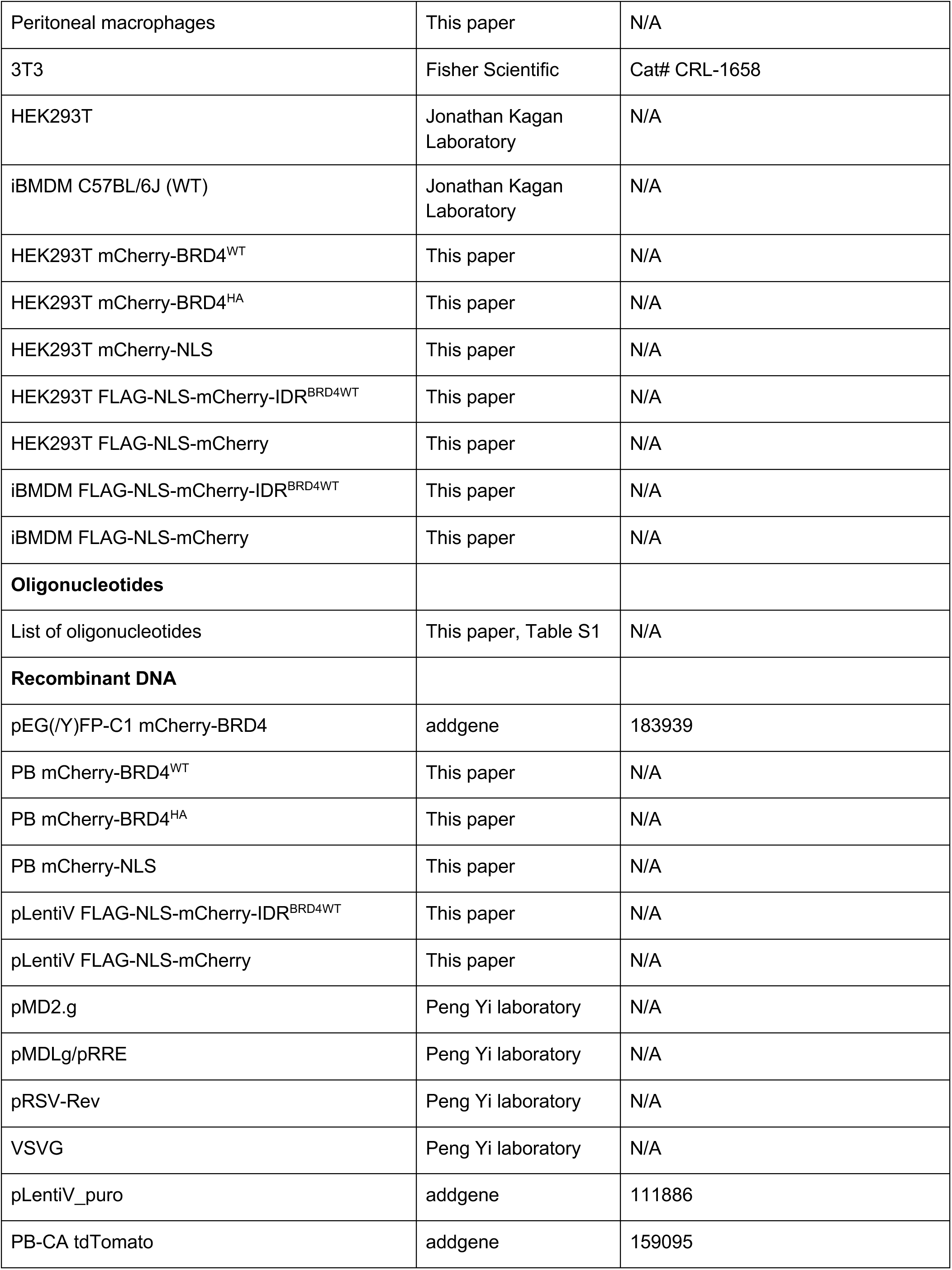
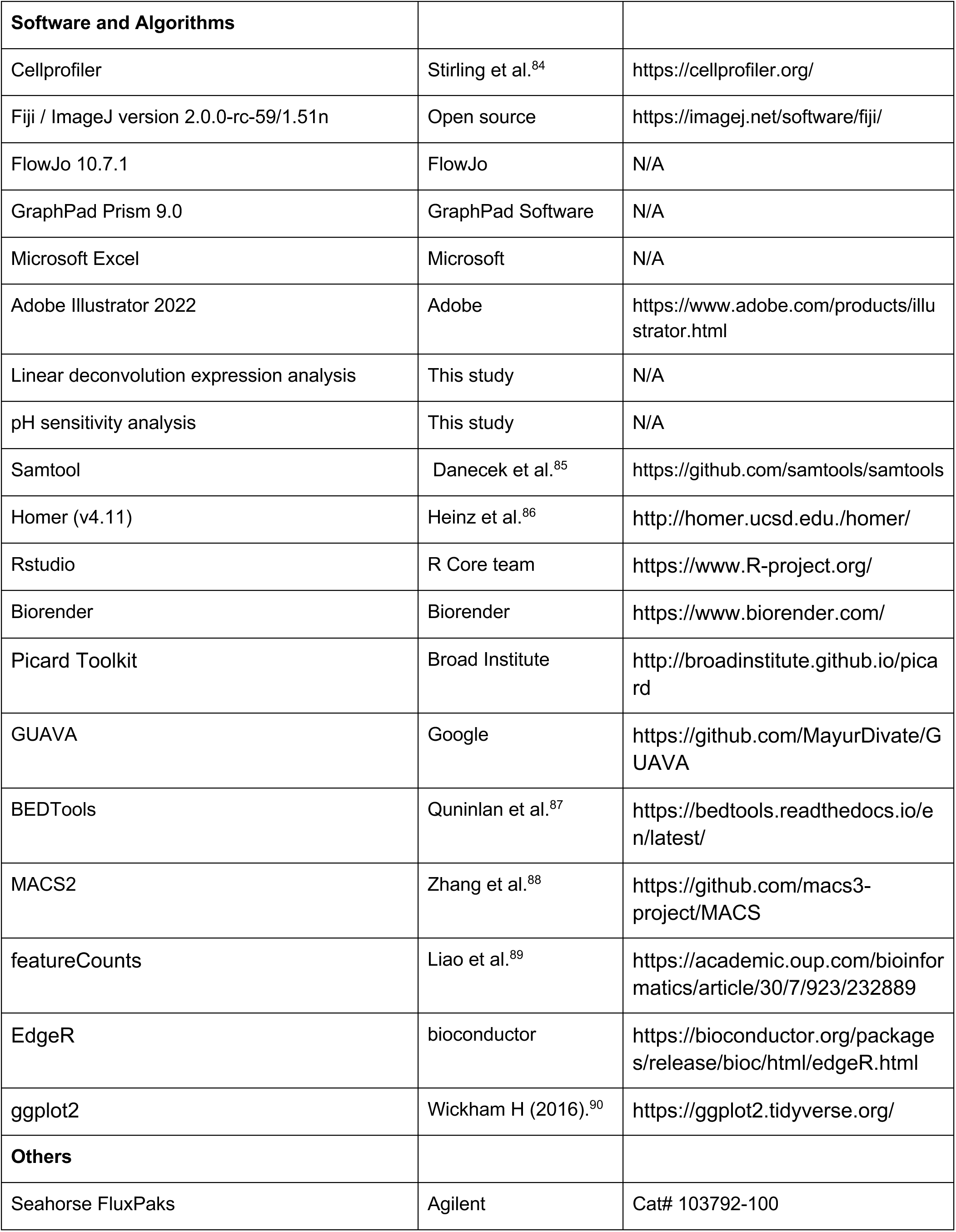
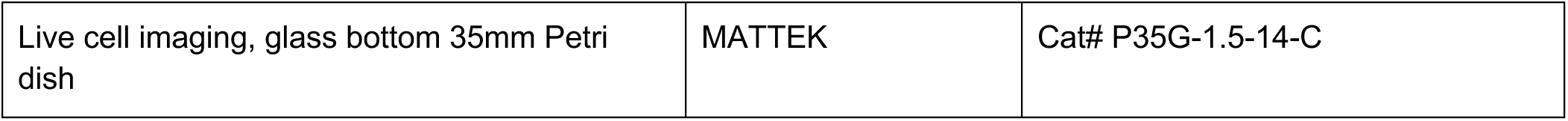

## RESOURCE AVAILABILITY

### Lead contact

Further information and requests for resources and reagents should be directed to and will be fulfilled by the lead contact, Xu Zhou

### Materials availability

All cell lines generated in this study will be available upon reasonable request from the lead contact.

### Data and code availability

All genomic data have been deposited on the NCBI Gene Expression Omnibus, with accession numbers GSE269299, GSE269891, GSE269298 and GSE269890. Original code created in this study will be available upon reasonable request. Any additional information required to reanalyze the data reported in this paper is available from the lead contact upon request.

## EXPERIMENTAL MODELS AND STUDY PARTICIPANT DETAILS

### Mice

Male and female C57BL/6J were bred and housed under specific pathogen free (SPF) facilities. All animal experiments were approved by Yale University or Boston Children’s Hospital Animal Care and Use Committee. C57BL/6J wild type, Hif-1α^flox^, Hif-2α^flox^ and Lyz2-Cre mice were purchased from Jackson Laboratory. Gpr65^-/-^ and Gpr68^-/-^ were purchased from KOMP repository and were crossed with C57BL/6J for at least 5 generations. Macrophage-specific HIF1α-deficient or HIF2α-deficient mice were created by crossing Lyz2-Cre and Hif-1α^fl/fl^ or Hif-2α^fl/fl^ mice. All mice were maintained in a specific pathogen-free facility and animal experimentation was conducted in accordance with institutional guidelines.

### Cell culturing

All cell lines were cultured at 37°C with 5% CO_2_. HEK293T, 3T3, iBMDM cell lines were grown in DMEM media (Gibco) supplemented with 10% FBS, 1x GlutaMAX (Gibco), 100 U/mL Penicillin-Streptomycin (Gibco), 1 mM Sodium Pyruvate (Gibco), and 10 mM HEPES (Gibco).

Bone marrow-derived macrophages are differentiated from bone marrow cells and cultured in macrophage growth medium (MGM) (RPMI + 2 mM L-glutamine, 1 mM sodium pyruvate, 10 mM HEPES, 200 U/mL penicillin/streptomycin, 10% FBS, 50 µM 2-mercaptoethanol and 30% L929-conditioned media).

### Preparation of Bone marrow-derived macrophages

The fresh isolated bone marrow cells from femurs and tibias were resuspended in macrophage growth media (MGM). The nonadherent cells were harvested next day (Day 1) and replated in 15 cm non-TC treated Petri dishes at 37°C in a 5% CO2 incubator. Cells were refed on day 4 with another 15 ml fresh MGM. Cells were collected with 3 mM ice-cold PBS-EDTA on day 6-7.

### Preparation of thioglycolate-induced peritoneal macrophages

Mice were injected with 1 mL autoclaved 3% brewer thioglycolate broth intraperitoneally. After 4 days, mice were injected intraperitoneally with 2 mg/kg LPS or PBS for 24 hours. The peritoneal macrophages were collected from peritoneal cavity with 6-9 mL cold PBS. The cells were immediately analyzed by flow cytometry for measuring intracellular pH. For imaging, cells were immediately plated onto 1% gelatin (Sigma) coated glass coverslips for immunofluorescence analysis.

### Preparation of human peripheral blood mononuclear cells

Human blood samples were collected from two healthy donors under the Boston Children’s Hospital Normal Blood Donor Services program. PBMCs were purified from whole blood by Ficoll gradient centrifugation. Briefly, whole blood was diluted 1:3 in RPMI-1640, and 30 mL of diluted blood was slowly layered over 15 mL of Ficoll-Paque (GE Healthcare, Piscataway, NJ), and centrifuged at 800 x g for 30 minutes at room temperature. PBMCs were collected, washed in RPMI-1640, and plated in tissue culture dishes for 24 hours to remove adherent cells. Next day, the floating cells were collected and seeded at 1 million cells in a 12-well plate with glass coverslip. Cells are deafferented in RPMI supplemented with 2 mM L-glutamine, 1 mM sodium pyruvate, 10 mM HEPES, 200 U/mL penicillin/streptomycin, 10% FBS, 50 µM 2-mercaptoethanol, and 50 ng/mL recombinant human CSF1. After 4 days, cells were refed with fresh growth medium. On day 7-10, the differentiated macrophages were used for experiments.

### Cell lines, Transfection, and lentiviral Transduction

For PB-mCherry-*Brd4*^WT^, the N-terminally mCherry-tagged full-length *Brd4* (mCherry-Brd4) was directly amplified from pEG(/Y) FP-C1 mCherry-Brd4 (Addgene #183939). To generate and PB-mCherry-Brd4^HA^, where 33 histidine residues within the IDR region (700-1400) were replaced with alanine, we employed a fragment assembly method. First, the coding sequence for histidine->Alanine BRD4-IDR mutant (IDR^HA^) was synthesized by Genscript. We acquired full length *Brd4^HA^* through overlapping PCR between the IDR^HA^ and *Brd4*^WT^, which was further amplified with mCherry-*Brd4*^WT^ to produce mCherry-*Brd4*^HA^. Both mCherry-*Brd4*^WT^ and mCherry-*Brd4*^HA^ were purified and cloned into a *piggyBac* backbone (Addgene #159095) using NEBuilder HiFi DNA assembly. To construct FLAG-tagged Brd4 constructs, FLAG-NLS was synthesized along with an overhang sequence matching the 5’ of mCherry. FLAG-NLS was assembled with mCherry-*Brd4*^WT^ or mCherry-*Brd4*^HA^ into a lenti-viral expression vector (Addgene #111886) using NEBuilder HiFi DNA assembly. We constructed Lenti-FLAG-NLS-mCherry-IDR^BRD4WT^ and Lenti-FLAG-NLS-mCherry vectors.

To generate transgene cell lines in 293T and iBMDM, *piggyBac* vectors containing the genes of interest were co-transfected with Pbase vector into 293T cells using lipofectamine 3000. Stable lines were enriched based on fluorescent signals. For transgene expression of BRD4 variants in macrophages, we utilized lentiviral system. Briefly, lentivirus carrying gene of interest were generated by co-transfecting lentiviral expression vectors and lentiviral packaging plasmids (pspax1, pmd2.g) into 293T cells using lipofectamine 3000. Viral supernatants were collected at post-transfection 48 hour and 72 hours, filtered, and applied to iBMDMs using spin-fection in the presence of Polybrene (10 μg/mL) at 1250 g for 60 min at 30°C. All cell lines were sorted based on mCherry signal to ensure comparable levels of transgene expression.

### Cell treatments

Acidic pH: pH medium was prepared using special RPMI-1640 without sodium bicarbonate. The media pH was adjusted to desired pH value using HCl (1 M) or NaOH (1 M). The cells were incubated at the indicated pH for at least 4 hours if without specification.

Inhibitors treatments: 500 nM JQ1, 50 nM MS645 and 2 μM iBET were applied 4 hours prior to LPS stimulation. 200 ng/mL cycloheximide, 100 mM Dibutyryl-cAMP, 5 mM Camptothecin, 50 mM TSA or 10 mM C646 was applied 1 hour prior to LPS stimulation.

1,6-hexanediol treatment: for immunofluorescent samples, 10% 1,6-hexanediol was added to cells for 2 min before fixation. For analysis of gene expression, we titrated down the concentration of 1,6-hexanediol so that cells remain normal morphology. We applied 1% 1,6-hexanediol for 2 hours after 2 hours LPS stimulation.

Conditioning cells for imaging at different pH: cells were incubated with medium at the indicated pH for 3.5 hours prior to treatment with 100 μM 2,4-Dinitrophenol solution for 30 minutes.

Manipulating intracellular pH after LPS stimulation: cells were treated with 10 μM nigericin and 120 mM KCl in pH 6.5 medium for 5 min, followed by pH 7.4 PBS wash for three times, and incubation at normal medium at pH 7.6 for 2 hours.

### H3k27Ac, H3k4me3, p65 and IRF3 ChIP-seq sample preparation and analysis

For ChIP-seq of histone modifications, p65 and IRF3, the experiments were performed as described.^91^ Briefly, mature BMDMs were plated at 15 million cells in 150 mm TC-treated plates. Cells were conditioned with pH 7.4 or 6.5 growth medium for 4 hours and stimulated with 10 ng/mL LPS for another 4 hours. Cells were crosslinked with 200 mM DSG-PBS for 30 minutes with continuous mixing, followed by 1% formaldehyde for 10 minutes, and quenched with 0.125 M glycine for 5 minutes. The chromatin from approximately 10 million cells (25-30 μg) were used for histone modification ChIP and 30 million cells (∼100 μg) were used for the transcription factor ChIP. Cell pellets were sonicated to an average chromatin fragment of 300-400 bp. The following antibodies were used for each ChIP experiment based on instructions: 10 μg anti-H3K4me3 (ab4729), 10 μg anti-H3K27Ac (ab8580), 1:100 anti-IRF3 (D83B9) and 10 μg anti-p65 (sc-372). 100μL of Protein G dynabeads (10003D) were used for each ChIP. Sequencing libraries were constructed using NEB Next Ultra II DNA Library Prep Kit for Illumina (E7645L).

Reads from Illumina paired end fastq files were mapped to the mouse genome (mm10) using Bowtie2 with the command “bowtie2-q--end-to-end--very-sensitive” to generate SAM files. Read duplicates were removed and BAM files were generated with the Picard toolkit. Enriched atac regions were identified using MACS2 using the options “’-f BAMPE--bw 200-B-g mm”. Bigwig files were generated using deeptools software using the bamCoverage tool with the options “--binSize 10--normalizeUsing RPGC--effectiveGenomeSize 2864785220--extendReads”. Bigwig files were visualized with UCSC genome browser as well as used to generate heatmaps (described below).

### BRD4 and BRD9 ChIP-seq sample preparation and analysis

For BRD4 and BRD9, ChIP-seq samples were prepared as previously described.^68^ 10-15 million BMDMs were crosslinked in 3 mM disuccinimidyl glutarate for 30 min then in 1% formaldehyde for 10 min at room temperature. After quenching with 125 mM glycine, the cells were washed in 1× PBS, pelleted, flash frozen in liquid nitrogen, and stored at 80°C. Cell pellets were thawed on ice and incubated in lysis solution (50 mM Hepes KOH pH 8, 140 mM NaCl, 1 mM ethylenediamine tetra-acetic acid (EDTA), 10% glycerol, 0.5% Nonidet P-40, and 0.25% Triton X-100) for 10 min. The isolated nuclei were washed (10 mM TrisHCl pH 8, 1 mM EDTA, 0.5 mM EGTA, and 200 mM NaCl) and sheared in (0.1% sodium dodecyl sulfate (SDS), 1 mM EDTA, and 10 mM Tris-HCl pH 8) with the Covaris E229 sonicator for 10 min. After centrifugation, chromatin was immunoprecipitated overnight at 4°C with 1:100 anti-BRD9 (Bethyl, pAb #A700-153) or 5 μg anti-BRD4 (Bethyl, Rabbit pAb #A301-985A50).The next day, the antibody-bound DNA was incubated with Protein A+G Dynabeads (Invitrogen) in ChIP buffer (50 mM Hepes KOH pH 7.5, 300 mM NaCl, 1 mM EDTA, 1% Triton X-100, 0.1% sodium deoxycholate, and 0.1% sodium dodecyl sulfate), washed, and treated with Proteinase K and RNase A and reverse cross linked. Purified ChIP DNA was used for library generation (NEBNext Ultra II DNA Library Prep Kit for Illumina) according to manufacturer’s instructions.

Single-end 100bp were aligned to the mouse genome mm10 using STAR alignment tool (version 2.5). ChIP-seq peaks were called using findPeaks within HOMER using-style factor (default settings, fourfold over local tag counts with FDR 0.001 [Benjamin–Hochberg]). Differential ChIP peaks were called using getDiffExpression.pl with fold change ≥ 1.5 or ≤ 1.5, Poisson P value < 0.0001. ChIP-seq peaks were annotated by mapping to the nearest transcription start site (TSS) using the annotatePeaks.pl program. Bigwig files were generated using makeUCSCfile on HOMER with flag-style chipseq and were visualized on UCSC genome browser.

### ATAC-seq analysis

BMDMs were plated on non-TC plates for ATAC-seq analysis. Cells were conditioned with pH 7.4 or 6.5 growth medium for 4 hours and stimulated with 10 ng/mL LPS for another 4 hours before harvesting with 3 mM ice-cold PBS-EDTA. Cell suspensions were counted and ∼ 50,000 cells were used for constructing each ATAC-seq library according to the protocol detailed by Buenrostro *et al*.^92^ ATAC-seq libraries were constructed using Illumina Tagment DNA Enzyme and Buffer Small kit.

Paired-end sequencing was performed with Next-seq 500 with paired end reads of 38 base pairs. Reads were then mapped to the mm10 mouse genome with Bowtie2 using the option-k 10-X 1000--end-to-end--very-sensitive”. Sam files were converted to Bam files, and read duplicates were removed using Picard. Blacklist regions and mitochondrial genomic reads were removed using BEDTools and Picard, respectively. Bam files were converted to Bed files also using BEDTools, and macs2 was used to call peaks using the converted bed file. Macs2 call-peak options used were “-f BED-g mm-q 0.05 – nomodel--shift-75--extsize 150--keep-dup all”. Macs2 peaks from each condition were concatenated and then overlapping regions merged using mergeBed from BEDTools. To count the number of reads in each merged peak, featureCounts was used. Read counts from featureCounts were normalized using the TMM method in EdgeR. Bigwig files were generated using using the bamCoverage (Deeptools) tool with the options--binSize 10-p max--normalizeUsing None--effectiveGenomeSize 2864785220--extendReads –scaleFactor X”. The scale factor “X” was derived from the TMM normalization. Bigwig files were visualized with UCSC genome browser as well as used to generate heatmaps (described below).

### Heatmap generation

To identify ATAC-seq peaks associated with each condition, the fold change between normalized read counts was quantified under each peak using R. Additionally, a peak was only considered to be associated with that condition if the merged Macs2 peak overlapped with the condition specific Macs2 peak. For whole gene plots, mm10 gene positions were downloaded from UCSC Genome Browser. Bed files from the condition specific ATAC-seq peaks and gene plots were quantified with the Bigwig files using the computeMatrix tool (deeptools) and visualized with the plotHeatmap tool (deeptools) to generate Heatmaps.

### RNA isolation and qRT-PCR

RNA was purified from cells using Qiagen RNeasy columns (Qiagen, 74106) according to the manufacturer’s instructions. cDNA was reverse-transcribed with MMLV reverse transcriptase using oligo-dT20 primers. qRT-PCR was performed on a CFX384 Real-Time System (Bio-Rad) using PerfeCTa SYBR Green SuperMix. Relative expression units were calculated as transcript levels of target genes over 1/1000 of Actb. Primers used for qRT-PCR are listed in Table S1.

### RNA-seq sample preparation and analysis

BMDMs were washed twice with ice-cold PBS and collected directly with RLT buffer (Qiagen RNeasy kit). RNA was purified from cells using Qiagen RNeasy columns with on-column DNase digestion according to the manufacturer’s instructions. Sequencing libraries were constructed following Illumina Tru-seq stranded mRNA protocol or NEB Next Ultra II stranded mRNA protocol. Paired-end sequencing was performed with Next-seq 500 with either 76 or 38 bp from each end.

Illumina fastq files were aligned with Kallisto program with default settings (Bray et al., 2016) in reference to mouse transcriptome annotation GRCm38 (ftp://ftp.ensembl.org/pub/release-90/fasta/mus_musculus/cdna/). The ENSEMBL IDs of each cDNA transcript were matched to the official gene symbols through BioaRt in R. The expression level for each transcript was reported in either TPM (transcripts per million) or as normalized counts, adjusted to a total of 10 million counts. For genes associated with multiple transcripts, the gene’s total expression was derived by summing the TPM or count values for all corresponding transcripts.

### Differential expression analysis

To identify differentially expressed genes, we utilized the Kallisto-Sleuth pipeline with the default settings. This analysis returned 34475 annotated genes in total in the mouse genome. The gene expression is calculated as tpm (transcript per million) or read counts per gene. We used read counts per gene for downstream analysis, after normalizing the total read counts to 10 million for each sample. For each pair-wise comparison, pH 7.4 LPS vs pH 7.4, pH 6.5 LPS vs pH 6.5, pH 7.4 vs pH 6.5, and pH 6.5 LPS vs pH 7.4 LPS, we calculated expression fold changes (with a pseudo count 2) and adjusted p-values with Sleuth. Using a stringent threshold (an adjusted p-value < 0.05, absolute fold change > 3-folds, and an average expression > 5 counts), we identified 1907 genes significantly regulated between any pair of tested conditions.

### Expression deconvolution analysis

To estimate the contribution of LPS stimulation, acidic pH and the interactions between these two conditions to the inflammatory response, we adopted the previously developed computational deconvolution algorithm.^54^ Briefly, when the contributions to gene expression are described quantitatively as fold activation, each experimental comparison listed above can be described as the sum of the following expression components in logarithmic scale. The expression components include LPS (the influence of LPS stimulation alone), pH (the influence of acidic pH alone), INT (the influence of the integration between LPS and acidic pH). Each of the comparisons in Figure S2B can be formulated as:

pH 7.4 LPS vs pH 7.4 = LPS

pH 6.5 vs pH 7.4 = pH

pH 6.5 LPS vs pH 6.5 = LPS + INT

pH 6.5 LPS vs pH 7.4 LPS = pH + INT pH 6.5 LPS vs pH 7.4 = LPS + pH + INT

The above formula can be transformed into matrix multiplication in the form of ***Y*** = ***X***\****β*** + ***ε***.

For each gene, ***Y*** represents the fold change matrix calculated from all pair-wise comparisons, ***X*** represents the deconvolution matrix, β represents expression components, and ε represents noise or error from experimental measurement. Linear regression analysis can be performed to infer the expression components while minimizing ε. Two biological replicates (each with a set of 4 RNA-seq data prepared with BMDMs from different source animals) were analyzed together by replicating the deconvolution matrix. The statistical significance is calculated for each β using the formula:

|β− β**_0_**| / **0** ∼ *t*-distribution with (***n***-***p***) degree of freedom. ***n*** is the number of measurement (10 with 2 sets of biological replicates) and ***p*** is the size of β (in this case 3). 0^2^ is the variance of the linear fit for each gene. To increase the specificity of the detection, we used β_0_ = 1.5 folds for positive expression component and β_0_ = −1.5 folds for negative expression component as the null hypothesis.

Among 1907 significantly regulated genes, we found 1620 genes containing at least 1 significant expression component of LPS, pH or INT (β>= 2-fold, p < 0.05, R^2^> 0.8). None of these genes include a significant error (ε). Majority of the expression variations measured between experimental conditions are well explained using the linear model with 3 expression components (Figure 2C). All 1620 genes are grouped based on whether a gene contains a significant LPS, pH or INT and the sign of expression components (Figure S2D). This generated 20 groups with each group containing at least 5 genes.

357 genes (Group3,4,6) were induced by LPS independent of pH. 171 genes (Group 4,7,8) were repressed by LPS independent of pH. Among pH-dependent LPS responses, 433 LPS-induced genes (Group 9,10,19) are repressed by acidic pH, while 58 genes are significantly enhanced (Group 15,16,17). Among pH-dependent LPS-dependent repression, the down-regulation of 272 genes (Group 12,13,20) was ameliorated at acidic pH, and 41 genes (Group 11, 18) showed stronger gene repression. Particularly, genes such as *Bcl3* (Group 5) and *Ccl3* (Group 6), are regulated by acidic pH independent of LPS stimulation. Their LPS-induced activation is independent of pH conditions, and thus are clustered into the pH-insensitive group.

### Gene ontology analysis

Gene lists were analyzed with Homer for functional enrichment. The resulting gene ontology terms were selected to illustrate representative differences between different gene groups. Genes from specific processes were selected to further illustrate expression differences between groups. The radius of the circle corresponds to the log2(FC), the transparency illustrates statistical significance, and the color indicates whether the changes are due to activation or repression.

### Nucleus isolation for examining BRD4 condensates

BMDMs were harvested using 3 mM ice-cold PBS-EDTA and spun down to remove excessive buffer. Cell pellet was washed with hypotonic buffer (20 mM HEPES pH 7.5, 10 mM KCl, 1.5 mM MgCl_2_, 1 mM EDTA, 0.1 mM Na_3_VO_4_, 0.2% NP-40, 10% glycerol, protease inhibitor cocktail (Roche)) and immediately spun down for 10 seconds at 10,000 g. Cell pellets were lysed in two volumes of ice-cold hypotonic buffer by passing through a 26-guage needle 3-4 times using a syringe. A cloudy supernatant indicates lysing of cells. The cell lysate was centrifuged at 10,000 g for 8-10 seconds to remove cytosolic componnets. The nuclear pellet was resuspended in PBS buffer at pH 7.4 and 6.5 in the presence of containing protease inhibitor cocktail and incubate on ice for 30 min. These nuclei were further seeded into a 24-well plate with matrigel-coated glass coverslips by spinning the plate at 1000 g. The attached nuclei were then fixed in 4% paraformaldehyde for 15 min at RT.

### Coimmunoprecipitation and Immunoblotting

The immunoprecipitation was performed as described previously.^93^ Briefly, about 1 x 10^7^ cells of FLAG-mCherry-IDR^BRD4^ or FLAG-mCherry 293T were harvested and washed with cold PBS. The cells were resuspended with 5 volumes of lysis buffer (50 mM Tris pH 7.4, 150 mM NaCl, 0.5% Triton X-100, 10% glycerol, 1 mM DTT, 1 mM PMSF and 1/100 Proteinase inhibitor cocktail), followed by incubation at 4°C with rotation for 30 min. Cell lysates were centrifuged at 12,000 g for 15 min at 4°C to remove debris. Next, 20 μL pre-equilibrated FLAG M2 beads (Sigma) were added to the supernatant and incubated overnight with rotating at 4°C. The beads were separated equally and washed with the wash buffer (50 mM Tris, 200 mM NaCl, 0.3% Triton X-100, 10% glycerol) at pH 7.4 or pH 6.5 for 5 times. Proteins were eluted from the M2 beads by boiling in 2 x LDS buffer for 10 min. One-third of the elution was analyzed on Mini-protean TGX stain-free protein gel (BIO-RAD, 4568126). Protein was transferred onto activated PVDF membrane using trans-blot Turbo system (BIO-RAD), and then blocked using EveryBlot Blocking Buffer (BIO-RAD, 12010020) for 5 min at room temperature. Primary antibody was incubated in Solution 1(TOYOBO, NKB-201) at 4°C overnight at recommended dilution (anti-BRD9 1:2000 and anti-FLAG 1:3000), and secondary antibody (1:5000) was incubated in 5% milk in TBST at room temperature for 1 hour. Samples were washed 3x with TBST following each round of antibody incubation. Fluorescent images were developed using ChemiDoc^TM^ MP Imaging System (BIO-RAD) and analyzed by Image Lab Touch Software Version 3.0.1.14 (BIO-RAD).

### Live cell imaging and immunofluorescence

For live cell imaging, full-length mCherry-BRD4^WT^ and mCherry-BRD4^HA^ 293T cell lines were seeded onto matrigel-treated glass-bottom confocal petri dishes (Mattek Corporation P35G-1.5-14-C). The cells were conditioned in pH 7.4 medium for 4 hours, followed by pH 6.5 medium without phenol red, in the presence of 100 μM 2,4-DNP. Imaging was performed using the Airyscan detector on a ZEISS 880 microscope equipped with live-cell imaging module. Each Cell was imaged with Z-stack series to capture the entire nucleus, on a 37°C heated stage with 5% CO2 supplementation. Images were acquired with 63x oil objective. Airyscan images were processed using Fiji (ImageJ) and CellProfiler.^84^

For immunofluorescence, 293T cells were plated on matrigel-treated glass coverslips, which enhanced the adherence of 293T cells. Macrophages were seeded on 0.1% gelatin-coated glass coverslips. After conditioning with different medium and treatments, cells were fixed in 4% paraformaldehyde for 15 min at RT, permeabilized with 0.5% Triton X-100 (Sigma Aldrich, X100) in PBS for 5 min at RT, and blocked in PBS containing 5% BSA (Sigma) and 0.3% Triton X-100 for at least 45 min at RT. Cells were stained with primary antibodies: anti-BRD4 (1:100 dilution), anti-MED1 (1:200 dilution), anti-HNRNPU (1:100 dilution), anti-SRSF2 (1:100 dilution), anti-F4/80 (1:100 dilution) and anti-CD11b (1:100 dilution) in 1% BSA overnight at 4°C, followed by incubation with secondary antibodies (Alexa fluor anti-mouse 647 or 488 or 555, Alexa fluor anti-rabbit 647 or 488, 1:1000) in the dark for 1 hour at RT. The samples were then counterstained with 20 μm/ml Hoechst 33342 for 5 min at RT in the dark, mounted on glass slide using ProLong Glass mounting media. Glass slides were sealed with Fixo gum (Marabu) and store at 4°C. Images were acquired in a Z-stack format using a ZEISS 880 laser scanning confocal microscope with 63x oil objective. Puncta or median fluorescence intensity was analyzed using Fiji(ImageJ) and CellProfiler. Statistical analysis was performed in GraphPad Prism.

### Condensate and Colocalization analysis

Immunofluorescent images acquired with Airyscan were transformed to gray-scale Tiff images using FIJI. Nucleus staining, BRD4 or MED1 TIFF images were imported into the “Speckle Counting” pipeline from CellProfiler to determine the foci objects of BRD4 and MED1 (available at https://cellprofiler.org/examples). The thresholds used for determining foci objects were adjusted with visual inspection and the same threshold was applied across all images within the same experiment. Colocalization Analysis was performed using a customized pipeline based on CellProfiler, using CorrectIlluminationCaculate, MaskObjects, CovertObjectsToImage, MeasureObjectIntensity, FilterObjects and RelateObjects in CellProfiler. This analysis generated data the percentage of overlapped foci objects in each cell, which were subsequently analyzed by GraphPad Prism.

### siRNA-mediated depletion of Brd4

To reduce the influence of endogenous human BRD4, 293T cells were simultaneous transfected with two *Brd4*-specific siRNAs or a Scrambled Negative Control. siBrd4 #1 (IDT, hs.Ri.BRD4.13.2, #426567433), siBrd4 #2 (IDT, hs.Ri.BRD4.13.4, #426567436), and Negative control DsiRNA, 5 nmol (IDT, #51-01-19-09) were listed in table S1. In brief, cells were cultured in 24-well TC plates and transfected at 50% confluency with 100 pmol of siRNA using Lipofectamine RNAiMAX (Invitrogen) according to the manufacturer’s protocol. After 24 hours, cells were transferred into matrigel-treated 35 mm Petri dish with the optics of glass bottom for 12-20 hours for live cell imaging.

### pHi measurement

pHi was measured by treating cells with 500 nM pH-sensitive dye SNARF^TM^-4F 5-(and-6)-carboxylic acid (Thermo Fisher Scientific, #S23921) in 1% BSA in PBS at 37°C for 20 mins, followed by flow cytometry analysis. pHi was determined as the ratio of fluorescence intensities measured at two emission wavelengths (580/42 nm and 640/60 nm). To calibrate SNARF fluorescence, we employed a high-potassium buffer (39.6 mM NaCl, 120 mM KCl, 2.3 mM CaCl_2_, 1 mM MgCl_2_, 5 mM HEPES, 10 mM glucose), spanning pH values from 6.0 to 8.0, in the presence of 10 uM nigericin (Sigma), which facilitate the external potassium ions with internal protons to equilibrate extracellular and intracellular pH level. The data shown are the representative results from 3 independent experiments, with 3 biological replicates in each condition.

### Seahorse metabolic assay

Differentiated BMDMs were seeded into 96-well Seahorse tissue culture plates at a density of 50,000 cells/well in complete macrophage growth medium adjusted to pH 7.4. Cells were allowed to rest at room temperature within TC hood for one hour to allow even distribution within the well, followed by overnight incubation at 37°C. Next day, cells were pre-treated with JQ1 inhibitor (500 nM) or vehicle (DMSO) for 30 mins prior to treatment with LPS (100 ng/mL) or vehicle in MGM at pH 7.4 for 5-6 hours at 37°C.

Subsequently, the RPMI medium was removed, and cells were washed twice with warm Seahorse assay medium XF RPMI Medium pH 7.4. Cells were then allowed to equilibrate at 37°C for 45 mins without CO_2_. Oxygen consumption rates (OCR, in pmol/min) or extracellular acidification rate (ECAR, in mph/min) were recorded on a Seahorse Bioscience XF96 Extracellular Flux Analyzer. The following inhibitors were added sequentially: for Mito-stress test, oligomycin (1.5 μM), FCCP (1.5 μM), and rotenone/antimycin A (0.5 μM); for glycolysis stress test, glucose (10 mM), oligomycin (1.5 μM) and 2-deoxyglucose (50 mM). The Seahorse medium was supplemented according to the assay performed: for the glycolysis stress test, the medium contained 2 mM glutamine; for the mitochondrial stress test, the medium contained 2mM glutamine, 10mM glucose and 1mM sodium pyruvate.

### *In silico* analysis of pH sensitivity

We downloaded the mouse proteome from UniProt (50,961) to analyze for putative proteins that are sensitive to a pH shift from 7.4 to 6.5. As most amino acids have minimal difference in protonation state at this pH range, we evaluated the expected charge shift of peptide sequences by focusing on their histidine residues. We applied a gaussian filter spanning 20 residues (Full Width at Half Maximum = 5; 0 = 2.12) on the histidine distribution to identify regions locally enriched with pH-sensitive residues.

The peaks of the gaussian smoothed charge profile corresponds to the value of ΔCharge. We selected all peptide regions that contain ΔCharge > 1 within a window of 20 AA (18596 regions). For each region, we extended 50 amino acid residues to the N-terminal side and to the C-terminal side of the βCharge peak. We used the metapredict function^61^ to calculate the consensus disorder score of this extended peptide region (∼100 AA) to evaluate overall disordered features. To ease the selection, we considered an extended peptide region as disordered if either N-terminal or C-terminal side has a consensus disordered score > 0.3 (12168 regions).^61^ Within these extended regions, we calculated the enrichment of proline and glutamine by inferring a PQ-score, as the peak value of the overall PQ density by summing of the gaussian smoothed (the same filter as above) densities of proline or glutamine individually. Similarly, the higher score between N-and C-terminal side of the ΔCharge peak was determined as the PQ score of the region. Applying gaussian filters to the charge distribution, proline or glutamine distributions helps identify regions containing clusters of residues with the desired features.

### Quantification and statistical analysis

Software for experimental quantifications is named in the respective method details sections. Statistical analyses were performed using the two-tailed, unpaired, Student’s t-test unless otherwise specified in figure legends. The P values were represented as follows: ns, not significant, *p<0.05, **p<0.01, ***p<0.001, ****p<0.00001.

## Supplementary figures

**Figure S1.**
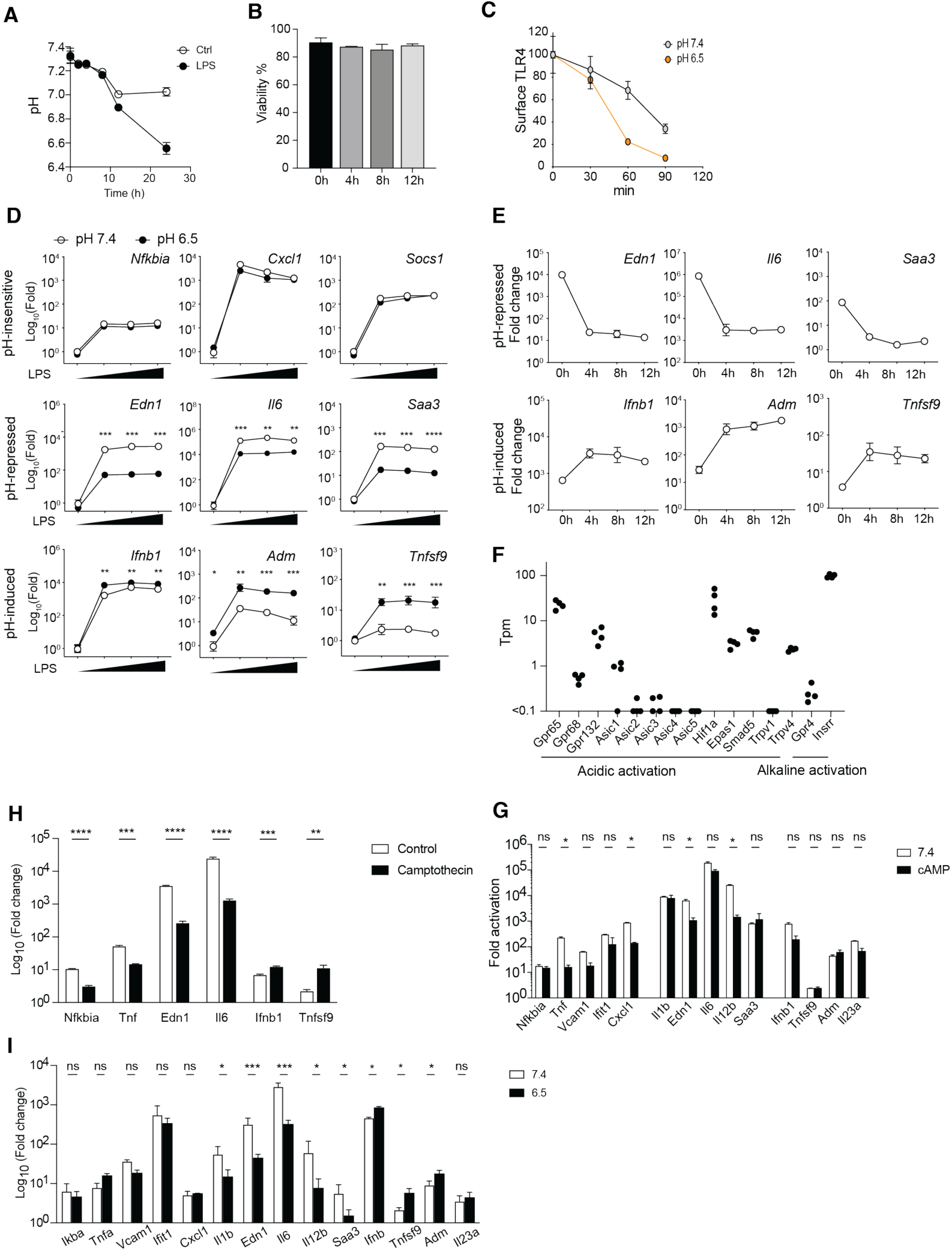
(**A**) Extracellular pH of BMDMs stimulated with 100 ng/mL LPS *in vitro*. Mean + STD. (**B**) Viability of BMDMs in acidic medium. Cell viability was assayed by flow cytometry using Annexin V and propidium iodide staining. Cells were gated for single cells based on FSC and SSC, and then gate for Annexin V^-^and PI^-^ for viable cells. (**C**) Internalization of TLR4 measured by flow cytometry. The median fluorescent intensity of each sample is subtracted of background (unstained control) and normalized to the surface TLR4 staining before LPS stimulation. Mean +/-STD. (**D**) Fold activation of inflammatory genes in BMDMs at 10-1000 ng/mL LPS treatment for 4 h. Unpaired t-test, Holm-Sidak’s test for multiple comparisons. mean +/-STD. (**E**) Fold activation of inflammatory genes in BMDMs after conditioning under acidic pH for 0-12 h. (**F**) Expression of known pH sensors in BMDMs from bulk RNA-seq. (**G-I**) Fold activation of inflammatory response genes after 4 h 10 ng/mL LPS, in the presence of 100 μM cAMP (G) or 5 μM Camptothecin (H) in WT BMDMs, or at pH 7.4 or 6.5 in *Nlrp3*^-/-^ BMDMs (I). Mean+/-STD. Unpaired Student’s t test, Holm-Sidak’s test for multiple comparisons. ns p>0.05, * p<0.05, ** p<0.01, *** p<0.001.

**Figure S2.**
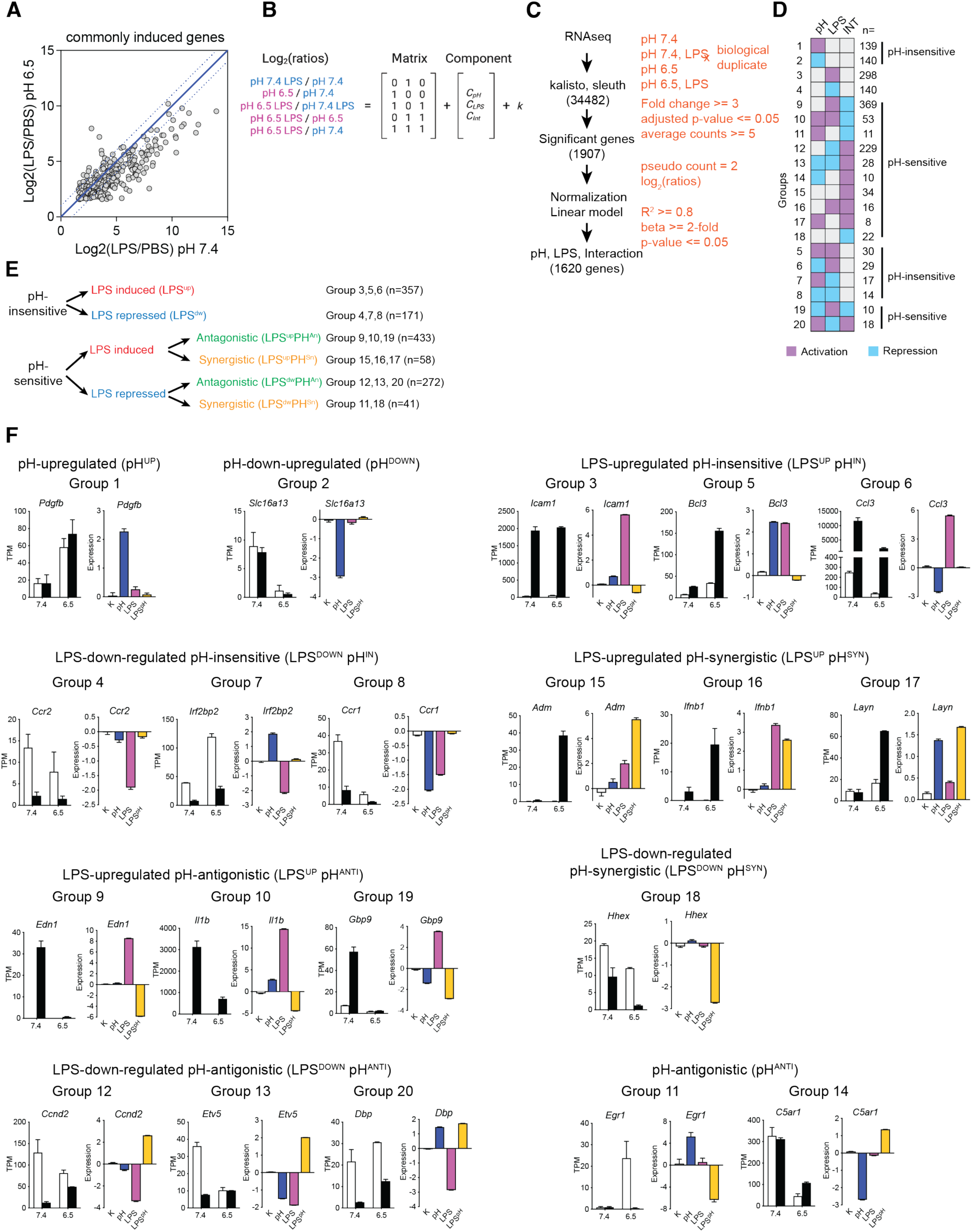
(**A**) Log2 (FC) of LPS-induced genes at both pH 7.4 and pH 6.5. Blue dot lines indicate the range of 2-fold variation. (**B**) Linear deconvolution matrix for dissecting the interactions between LPS and pH. (**C**) Analytic pipeline and thresholds applied to identify pH-and LPS-regulated genes. (**D**) Illustration of pH-sensitive and pH-insensitive genes in 20 identified clusters. (**E**) illustration of combining cluster groups into pH^IN^, pH^ANTI^, and pH^SYN^ groups. (**F**) Examples of each 20 gene clusters based on deconvoluted gene expression components.

**Figure S3.**
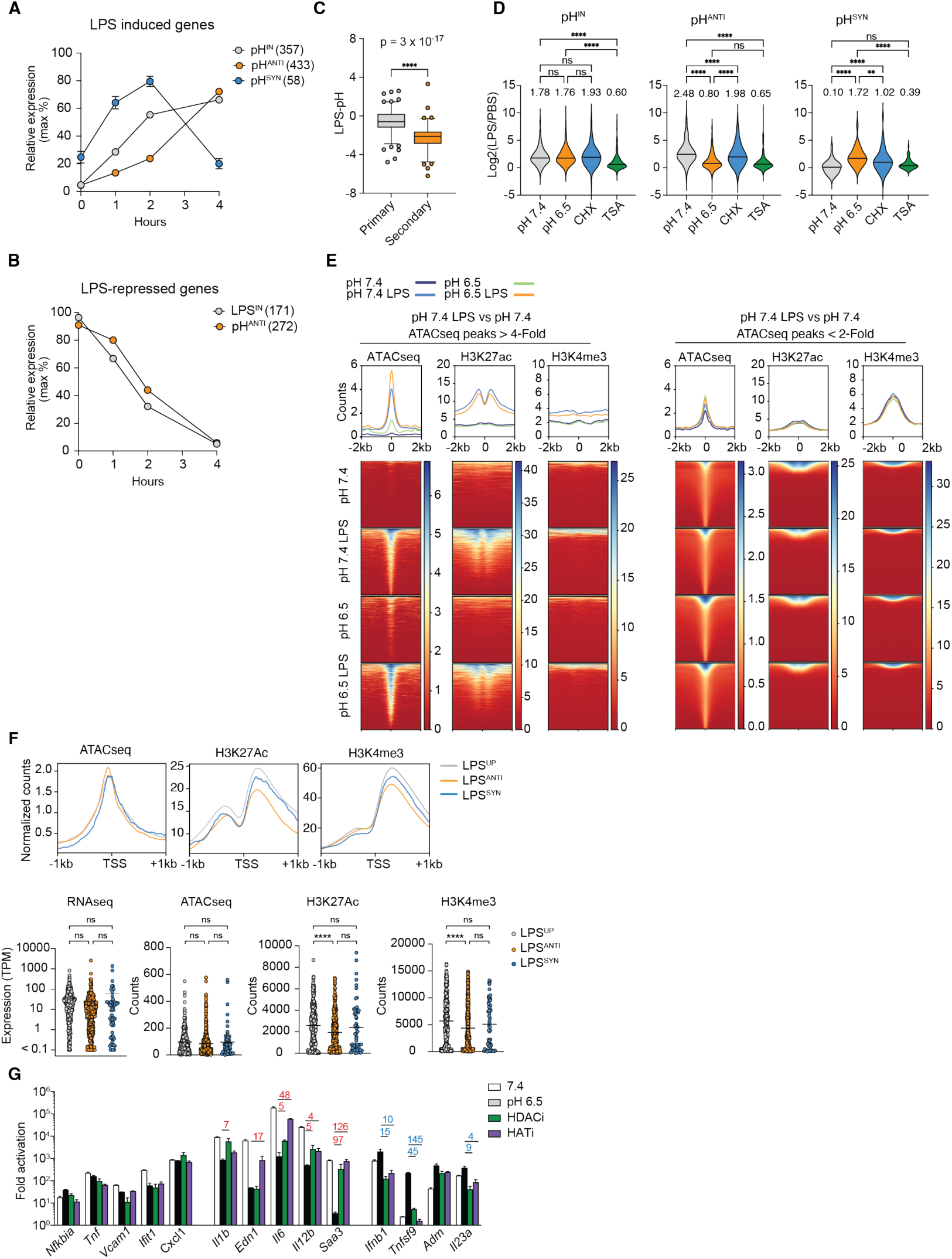
(**A**) Expression kinetics of all LPS-induced genes for pH-insensitive (357), pH-antagonistic (433) and pH-synergistic (58) groups. **(B**) Expression kinetics of all LPS-repressed genes for pH-insensitive (171) and pH-antagonistic (272) groups. (**C**) The LPS-PH component of primary and secondary response genes in response to LPS in BMDMs identified in Tong et al., 2016. (**D**) Violin plots of the fold change of gene expression between 10 ng/mL LPS at 4 hours and PBS control for the indicated conditions. CHX, 200 ng/mL cycloheximide; TSA, 50 mM TSA. The numbers above violin plots indicate the group median. Friedman ANOVA, Dunn’s test for multiple comparisons. (**E**) Genomic profile of ATAC-seq, ChIP-seq of H3K27Ac and H3K4me3 marks at pH 7.4, pH 7.4 LPS 4 h, pH 6.5, pH 6.5 LPS 4 h, for ATAC-seq peaks with significant increase after LPS at pH 7.4 (left), or with less than 2-fold change at pH 7.4 (right). (**F**) Average profile of ATAC-seq, ChIP-seq of H3K27Ac and H3K4me3 marks AT pH 7.4 for LPS^IN^, LPS^ANTI^ and LPS^SYN^ groups (top) and comparison of RNA-seq, ATAC-seq, H3K27Ac and H3K4me3 signals for for LPS^IN^, LPS^ANTI^ and LPS^SYN^ groups (bottom). (**G**) Activation of inflammatory genes by 10 ng/mL LPS for 4 h, with 50 mM TSA (HDAC inhibitor) or 10 μM C646 (HAT inhibitor). ns p>0.05, ** p<0.01, **** p<0.0001.

**Figure S4.**
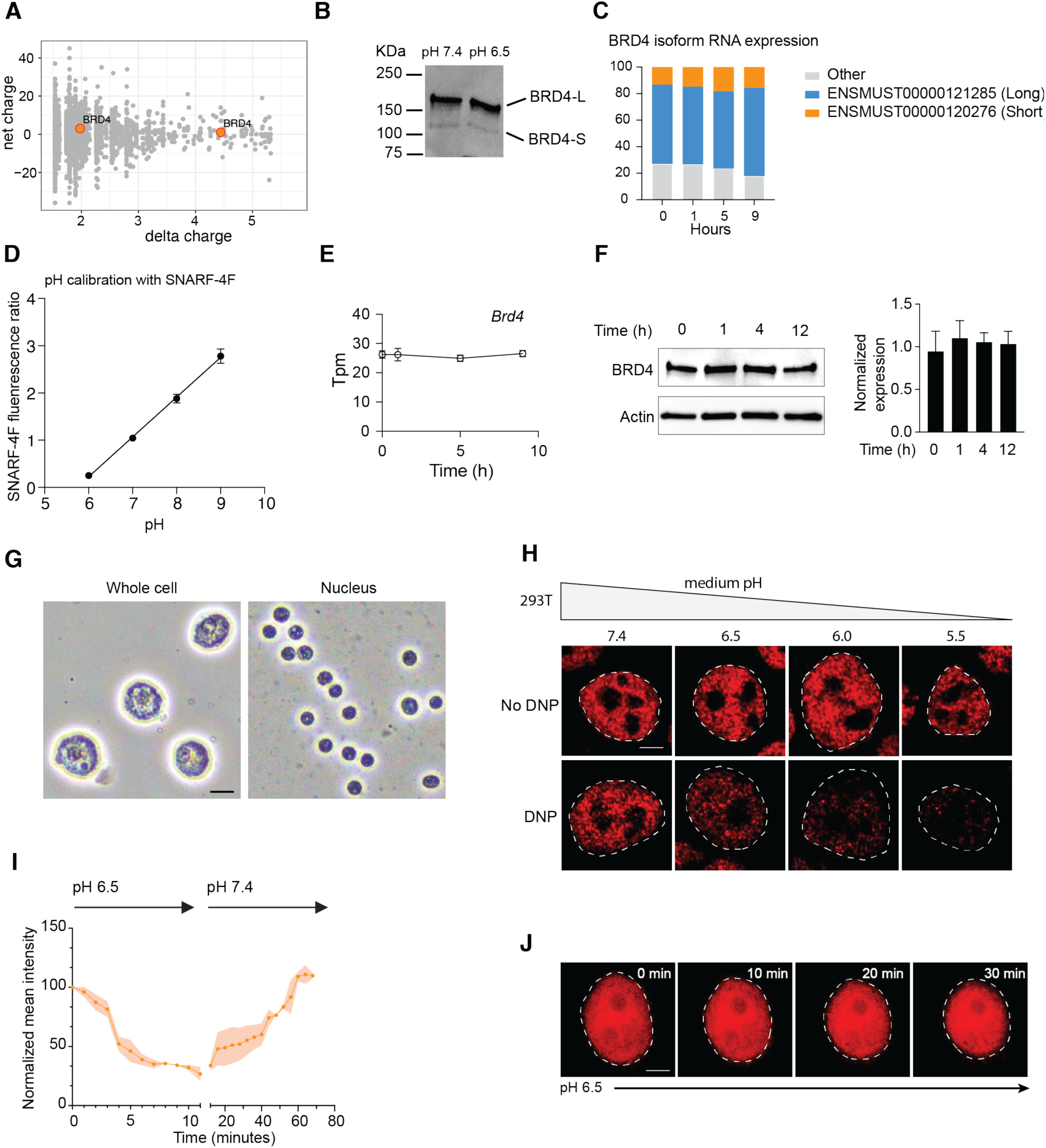
(A) Bioinformatic analysis of Δcharge and total charge of amino acid side chains for pH-sensitive peptide sequences shown in Figure 4A. **(B)** Western blot of BRD4 isoforms at pH 7.4 and pH 6.5. **(C)** Expression of various *Brd4* isoforms in murine BMDMs at pH 6.5. **(D)** Calibration curve for measuring pHi with SNARF-4F. R^2^=0.999. **(E)** Gene expression of *Brd4* in BMDMs at acidic pH. **(F)** Western blot of BRD4 proteins in BMDMs under acidic pH for various time points. **(G)** Imaging of whole BMDM cells and isolated nuclei from BMDMs. **(H)** Immunofluorescent staining of BRD4 in 293T cells (E) conditioned at pH 7.4 - pH 5.5 for 4 hours, with or without 100 mM 2,3-DNP for 0.5 h. (**I)** Quantification of BRD4 in 293T cells for live cell imaging in Fig.4G. **(J)** Time lapse imaging of 293T cells stably expressing mCherry at acidic pH. Scale bar represents 2 mm.

**Figure S5.**
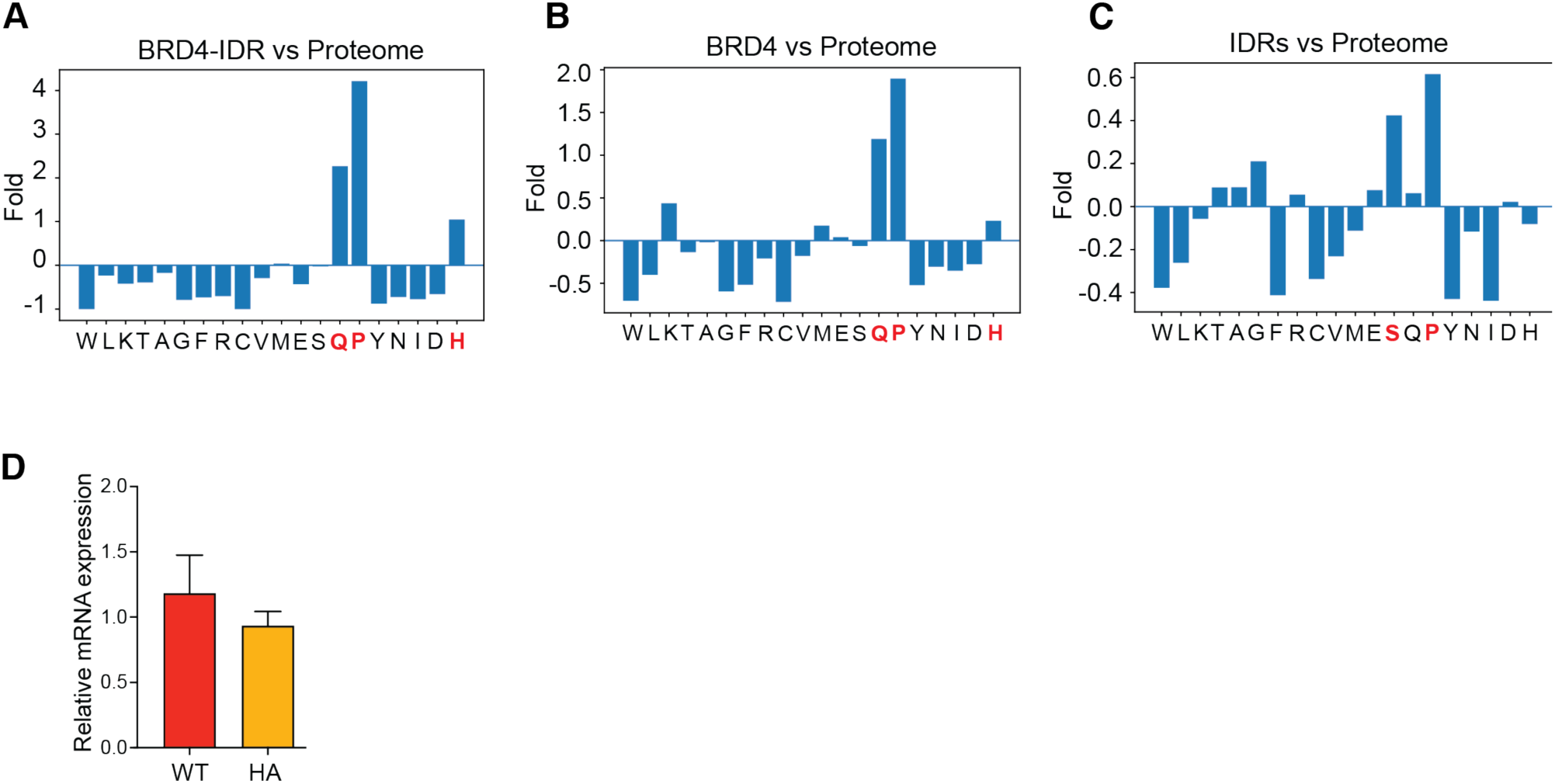
(**A-C**) Relative enrichment of amino acids between BRD4-IDR and the mouse proteome (A), BRD4 and the mouse proteome (B), and all IDRs and the mouse proteome (C). (**D**) Expression of exogenous BRD4 in BRD4^WT^ (WT) and BRD4^HA^ (HA) in 293T cells.

**Figure S6.**
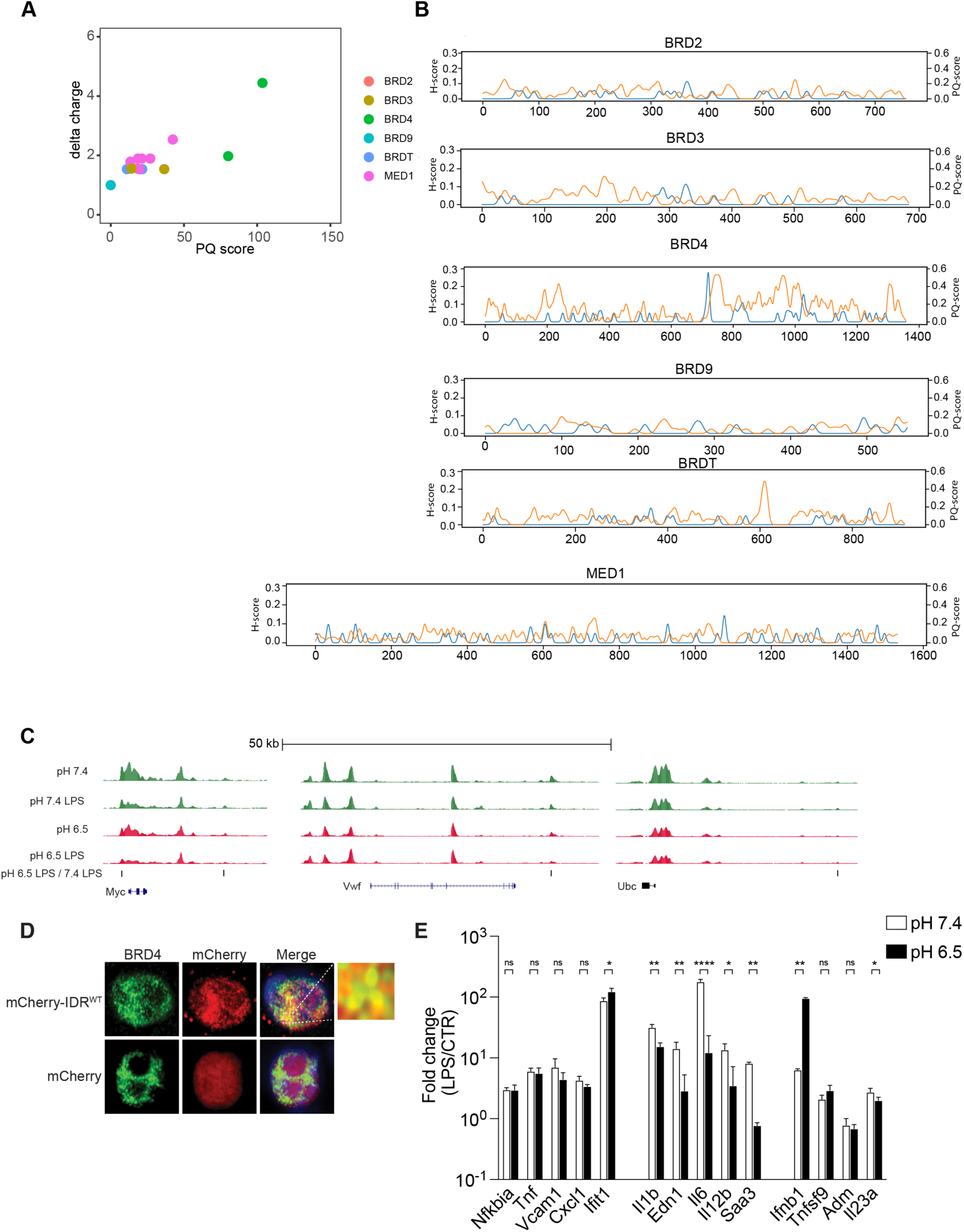
(**A**) Bioinformatic analysis of pH-sensitivity of BET family proteins and MED1. (**B**) Tracks of H, P and Q scores for full length BRD2, BRD3, BRD4, BRDT, BRD9 and MED1. (**C**) BRD4 ChIP-seq tracks at house-keeping genes *Myc*, *Vwf* and *Ubc*. Lines underneath the tracks mark the peaks with significant reduction of BRD4 occupancy comparing pH 6.5 LPS and pH 7.4 LPS. (**D**) Immunofluorescent imaging of mCherry fused BRD4-IDR (red), mCherry (red) and endogenous BRD4 (Green) in 293T cells. (**E**) Fold activation of inflammatory genes in iBMDMs after 6 h 100 ng/mL LPS at pH 7.4 or 6.5, normalized to unstimulated conditions respectively. Mean+/-standard deviation (STD). Unpaired t-test, Holm-Sidak’s test for multiple comparisons.

**Figure S7.**
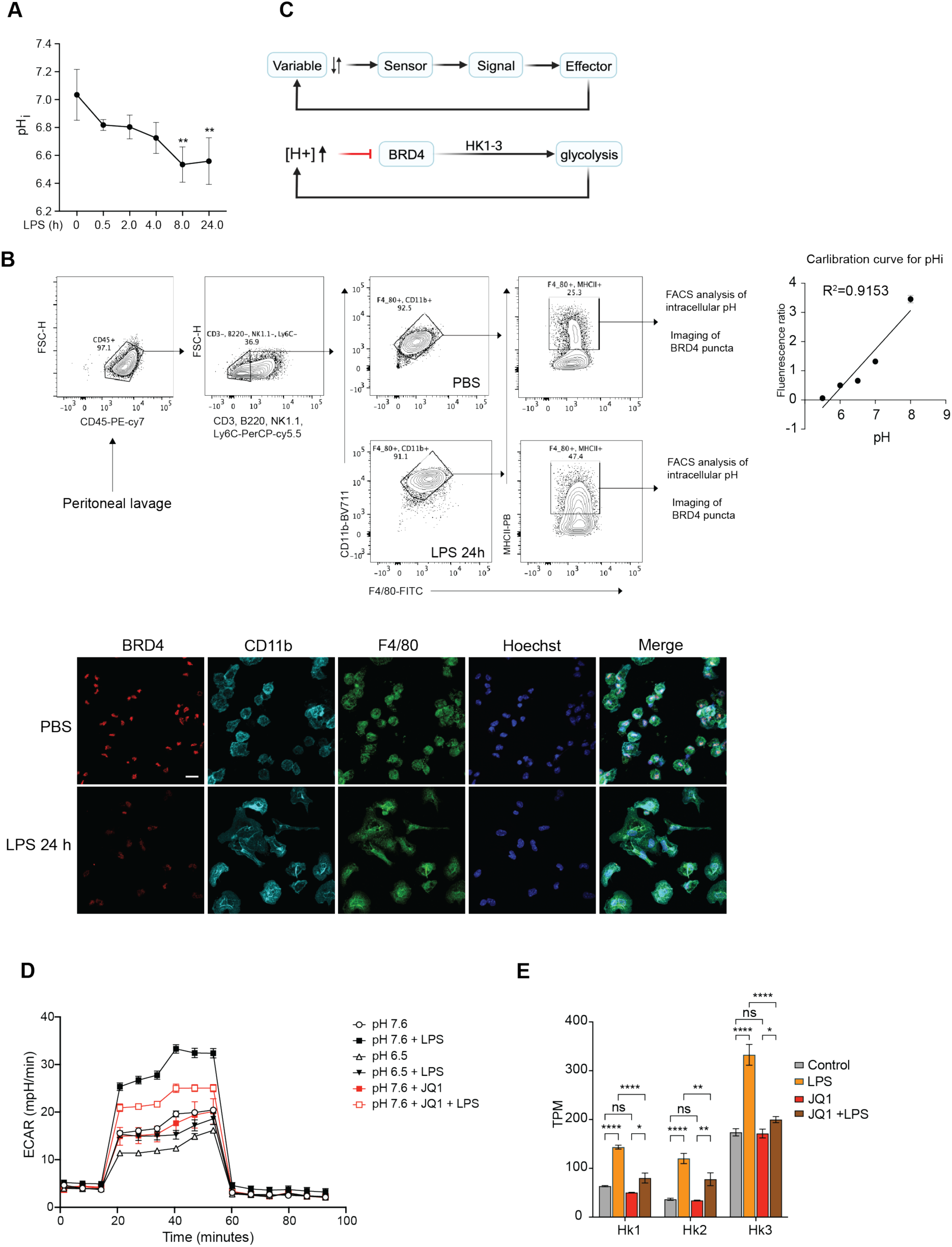
(**A**) pHi of BMDMs at various time points after 100 ng/mL LPS stimulation. ** p<0.01. (**B**) Flow cytometry gating strategy to isolate peritoneal macrophages, pH calibration, and immunofluorescent imaging of peritoneal macrophage staining 24 h after 3 mg/kg LPS treatment. (**C**) Diagram illustrating BRD4 as pH-sensor to control inflammatory response. (**D**) Seahorse analysis of ECAR on BMDMs treated with JQ1 or acidic pH. (**E**) The expression of *Hk1*, *Hk2* and *Hk3* with 100 ng/mL LPS or 0.5 μM JQ-1 treatment.

